# Annual community patterns in the *Halichondria panicea* sponge microbiome are characterized by seasonal switching between sponge-specific marine bacteria

**DOI:** 10.64898/2026.04.20.719716

**Authors:** Leon X. Steiner, Ute Hentschel

## Abstract

This study investigates the seasonal dynamics of the microbiome within the marine sponge *Halichondria panicea* from Baltic coastal waters, focusing on its symbiotic relationship with *Candidatus* Halichondribacter symbioticus. Over 16 months, we observed distinct summer and winter microbial communities, transitioning rapidly between these states during spring and fall. Marine sponges host complex microbiomes composed of diverse microbial taxa that play critical roles in host metabolism and nutrient cycling within marine ecosystems. While our understanding of sponge microbiomes has traditionally been based on static characterizations, the temporal dynamics of these associations across seasonal cycles remain poorly understood. In this study, we investigated temporal variation in bacterial symbionts of *Halichondria panicea* over 16 months in Baltic coastal waters using high-throughput amplicon sequencing of bacterial 16S rRNA gene sequences. The microbiota of *H. panicea* exhibited host-specific structure and a high degree of stability across seasons, despite fluctuations in environmental factors such as temperature, salinity, photoperiod intensity, and inorganic nutrient availability. In contrast, bacterial communities in surrounding seawater displayed large seasonal shifts which potentially mix with the sponge bacterial community, suggesting that different degrees of ecological pressures act on free-living and symbiotic marine bacteria. These findings establish an empirical baseline for identifying abnormal shifts in symbiont communities, which could be indicative of environmental stress or biological disturbance events.

## Introduction

Marine sponges are ancient metazoans that host uniquely diverse and complex microbial communities, forming mutualistic relationships crucial to their survival and ecological role. These microbiomes facilitate essential nutrient exchanges and produce chemical defenses, attributed by the sponges’ ability to thrive in diverse marine environments (Müller 2003; Wulff 2006; Laport, Santos, and Muricy 2009). The microbial composition varies between high microbial abundance (HMA) and low microbial abundance (LMA) sponges (Hentschel et al. 2003). HMA sponges, with dense and structurally complex tissues, support abundant and diverse microbial populations that play critical roles in nutrient cycles such as carbon fixation and nitrogen transformations. In contrast, LMA sponges, characterized by higher filtration rates, rely less on microbial symbionts for dissolved organic matter (DOM) processing and more on particulate organic matter (POM) consumption (Hentschel, Usher, and Taylor 2006; Hentschel et al. 2012). These distinctions highlight how sponge-microbe symbioses have evolved to adapt to varying ecological niches, enabling significant biogeochemical impacts in marine ecosystems.

Over time, marine sponge microbiomes exhibit changes ranging from subtle shifts in a few rare taxa to dramatic restructuring of entire communities, with variability depending on species, environmental conditions, and life stage (Glasl et al. 2018). At the mild end of the spectrum, some host-symbiont associations remain remarkably stable despite seasonal fluctuations. Elevated seawater temperatures significantly impacted the ecophysiology of *Geodia barretti*, contributing to the mass mortality events at Tisler Reef, Norway, but the microbiome remained stable (Strand et al. 2017). Additionally, three *Ircinia* species in the northwest Mediterranean Sea maintain consistent, host-specific bacterial communities over at least 1.5 years, even as temperature and irradiance vary (Pita et al. 2013). Although these stable holobionts show slight seasonal shifts among their less abundant symbionts, their core communities persist, demonstrating that minor disturbances do not necessarily topple established microbial partnerships.

In other species, subtle seasonal influences give way to more pronounced microbiome changes as environmental conditions become increasingly challenging (Yang, Zhang, and Franco 2019). *Mycale acerata* in Antarctica exhibits low variability despite extreme habitats, yet modest seasonal adjustments still occur (Happel et al. 2022; De Castro-Fernández et al. 2023). *Hymeniacidon sinapium* from the Yellow Sea, however, experiences more noticeable microbial rearrangements linked to temperature and sedimentation (Cao et al. 2012), while *Halichondria panicea* also displays clear temporal variations with less defined environmental drivers (Knobloch, Jóhannsson, and Marteinsson 2019; Schmittmann 2021; Al-Haddad, Caldwell, and Clare 2025). Here, the microbiome composition does not collapse, but these patterns indicate a holobiont more sensitive to environmental shifts than the stable *Ircinia* holobionts.

Further along this gradient, conditions such as altered salinity and anthropogenic impacts like aquaculture cultivation provoke stronger responses (Laroche et al. 2021). Under hyposaline conditions, *Coscinoderma matthewsi* experiences a marked reduction in *Chloroflexi* and an increase in *Proteobacteria*, signaling a community pushed toward more generalist taxa (Glasl et al. 2018). Similarly, sponges moved to mesocosm environments or aquaculture settings, such as *Mycale laxissima*, gain new microbial associates over time, often increasing in diversity but drifting away from their original stable symbiont assemblages (Mohamed et al. 2008). Disturbances in sponge microbiomes can manifest as changes in microbial diversity, with either increases or decreases signaling imbalance. For example, *Aplysina aerophoba* exhibited higher microbial diversity in diseased individuals, attributed to the proliferation of opportunistic pathogens and changes in bacterial group abundances (Monti et al. 2022). These shifts, though significant, often remain at least partially reversible if conditions return to normal, reflecting adaptability without complete breakdown. The most extreme alterations occur when sponges confront severe, prolonged stressors like sustained elevated temperatures. *Rhopaloeides odorabile*, when incubated at 32°C, loses key stable symbionts and becomes dominated by opportunistic microbes from *Proteobacteria* and *Bacteroidetes* (Simister et al. 2012; Fan et al. 2013). This constitutes a fundamental restructuring that can impair nutrient cycling or chemical defense, challenging the holobiont’s resilience and potentially impacting sponge health. Not all species recover swiftly from such radical changes.

Beyond external factors, development and reproduction also introduce microbiome variability. In *Amphimedon queenslandica* (Fieth et al. 2016), the microbiome reorganizes shortly after larval settlement, transitioning from vertically inherited symbionts to environmental recruits that stabilize as the sponge matures. Such ontogenetic shifts represent controlled restructuring, a necessary recalibration for the sponge’s changing physiology and ecological role. Even here, the extent of alteration is finely tuned, with early disturbances paving the way for a functionally stable adult community (Turon et al. 2024). Research into the mechanisms of microbial symbiont selection and maintenance in marine sponges has increased, with evidence suggesting that both stochastic processes and microbial transmission modes (vertical and horizontal) are involved. The concept of “leaky vertical transmission” or “mixed-mode transmission” was proposed to explain how sponges acquire their microbiomes (Oliveira et al. 2020). While vertical transmission has been observed in many sponge species, it does not fully account for the complete bacterial community. For instance, Mediterranean sponge larvae share only a small fraction of bacterial ASVs with their parents (Björk et al. 2019), and horizontal transmission has been observed to contribute significantly to the bacterial community, with sponge species often sharing core bacteria with the surrounding seawater (Oliveira et al. 2020). Despite these findings, the diversity and significance of these transmission modes remain largely unresolved.

Longitudinal microbiome studies, which track the same individuals over time, offer significant advantages in understanding microbiome dynamics by controlling inter-individual variability and enabling the detection of subtle temporal changes. Landmark studies of the human microbiome (Human Microbiome Project Consortium 2012; David et al. 2014), have demonstrated how repeated sampling reveals patterns of microbiome stability, resilience, and responses to perturbations, such as dietary shifts or environmental factors. These studies have established a gold standard and highlighted the critical need for temporal resolution in establishing causal relationships and identifying rare or transient microbial events, trajectories of microbiome development, and responses to stressors. Due to the inherent difficulty of working in fluctuating marine environments, most temporal microbiome studies in sponges have relied on random sampling and interspersed snapshots without tracking individuality.

This study focuses on the sponge species *Halichondria panicea*, commonly known as the breadcrumb sponge, widely distributed in temperate coastal waters (Goldstein and Funch 2022). As an LMA sponge, H*. panicea* hosts microbial communities at densities comparable to surrounding seawater but distinct in composition (Wichels et al. 2006), including both core symbionts and transient taxa. Previous research has provided valuable insights into the microbiome dynamics of *H. panicea*, revealing both temporal variability and host-specificity (Knobloch, Jóhannsson, and Marteinsson 2019; Al-Haddad, Caldwell, and Clare 2025). Time-series studies (Wichels et al. 2006; Lamb and Watts 2023) suggest that while the microbiome of sponges remains stable in response to seasonal temperature changes, temporal fluctuations can occur due to factors beyond predictable environmental patterns, such as stochastic events or host-specific traits. Habitat-related studies further indicate that local conditions, such as nutrient availability and water flow, may influence microbiome variability within and among individual sponges (Barthel 1991; F. Lüskow et al. 2019; Florian Lüskow et al. 2019), emphasizing the potential role of microhabitat in shaping microbial communities . These findings contribute to the hypothesis that LMA sponges, including H. panicea, may exhibit greater flexibility in their host-symbiont interactions compared to HMA sponges, allowing dynamic responses to environmental fluctuations. However, the mechanisms underlying this flexibility and its ecological implications remain poorly understood.

Applying these principles to marine sponges could similarly elucidate the stability and variability of sponge-microbe associations across seasons or environmental shifts. By investigating the temporal dynamics of the *H. panicea* microbiome, this study aims to build on existing knowledge, exploring how environmental and host-individual factors shape microbial diversity and stability. Understanding these dynamics is critical for elucidating the ecological roles of LMA sponges and their microbial symbionts in temperate marine ecosystems. This approach would pave the way for a deeper understanding of temporal dynamics in sponge holobionts, offering novel insights into their ecological resilience and responses to environmental changes.

## Materials and Methods

### Study site and sample collection

The study was conducted at a temperate coastal site in Kiel, Germany (54.424705 N, 10.175133 E), at a public beach with a wavebreaker. The site was chosen for its ease of access, close proximity to the research institute, and has been the focus of previous studies related to sponge symbioses of *Halichondria panicea*. Up to 15 stations around the wavebreaker were established by marking single sponge individuals and colonies for repeated sampling, and their GPS coordinates recorded. When a sponge or its marking was lost, another one from the same station replaced it on the next sampling event. When an entire station was lost, a new station at the same location or close proximity to it was established at the next sampling event.

Sponge samples were collected monthly, from February 2022 including May 2023, over 16 months from each station by SCUBA diving, ensuring consistent methodological conditions for repeated sampling of marked sponges. Collections consistently took place during morning hours (8:00-13:00 h), ensuring sampling of sponge states during larval release in spring which peaks in the morning. Sponge tissue biopsy samples (∼3 cm^3^) were cut underwater with scalpels, minimizing surrounding damage to ensure sponge survival, and collected with ambient seawater in 50 mL tubes. Additionally, 20 L of seawater was sampled from ∼1 m depth near the sponge colonies. All samples were stored on ice during transportation, and were further processed at the institute within the same day.

At the lab, sponge tissue samples were repeatedly washed with sterile artificial seawater of corresponding practical salinity. Contaminating tissues and organisms (e.g. brown and red algae, mussels, copepods) were removed by dissection, clean tissues chopped into small pieces, and stored at -80°C. Batches of seawater were processed in parallel and by a series of filtration steps. Removal of particulate matter and crude contaminants was done with a 20-25 µm prefiltration step through Miracloth (Merck KGaA, Darmstadt, Germany), followed by vacuum filtration with Nalgene bottle top filters (Fisher Scientific GmbH, Schwerte, Germany) and a 5 µm PVDF filter (Merck KGaA, Darmstadt, Germany). To collect the bacterial fraction, the seawater filtrate was further filtered with a similar clean setup on a 0.22 µm PVDF filter (Merck KGaA, Darmstadt, Germany). Finally, under a clean bench, the 0.22 µm filter was removed, folded into itself, and stored at -80°C as well.

### Environmental data

On site, a HOBO Pendant Temperature/Light 64K Data Logger (HOBO Data Loggers, MA, USA) was deployed underwater next to the sponge colonies at a depth of ∼1.5 m, to measure hourly seawater temperature and subsurface light intensity parameters. The logger operated from March 2022 until March 2023. Additional daily environmental seawater parameters for the Kiel Fjord were taken from the KIMOCC environmental data monitoring, at the GEOMAR pier (54.330192 N, 10.149911 E) at ∼1 m depth, from Feb 2022 and May 2023 regarding seawater temperature and practical salinity (Hiebenthal, Begler, and Melzner 2023). Further monthly measurements regarding seawater temperature, practical salinity, oxygen saturation, and nutrient levels in terms of nitrite (NO_2_), nitrate (NO_3_), phosphate (PO_4_), and silicate were retrieved from the Boknis Eck Time Series Station (54.513505 N, 10.022310 E) from Feb 2022 to May 2023 (Lennartz et al. 2014), as well as the Kiel Lighthouse (54.5 N, 10.266667 E) from the Federal Maritime and Hydrographic Agency (http://www.bsh.de). Additional daily nutrient measurements from February 2022 to April 2023 were reverse engineered from the GEOMAR pier time series data (54.330083 N, 10.150528 E) published elsewhere (Kim 2024). Seasonal averages were computed from all available daily and monthly measurements, and further missing monthly averages were interpolated with the {forecast} package in R (Hyndman and Khandakar 2008) for correlation analyses.

### DNA extraction and sequencing

DNA was extracted from <100 mg sponge tissue and half a circular filter for seawater samples with the DNeasy PowerSoil Kit (Qiagen, Netherlands). The DNA was eluted in 50 μL elution buffer, quantified by Qubit (DNA BR and HS Kits, Thermo Fisher Scientific, USA) and purity checked with NanoDrop 2000c (Thermo Fisher Scientific, USA) (Schmittmann and Pita 2022). The V3-V4 variable regions of the 16S rRNA gene were amplified in a one-step PCR using the primer pair 341F-806R (dual-barcoding approach (Kozich et al. 2013); primer sequences: 5’-CCTACGGGAGG-CAGCAG-30 and 5’-GGACTACHVGGGTWTCTAAT-30). Paired-end sequencing (2×300bp) was conducted on the MiSeq platform (Illumina, San Diego, USA) with v3 chemistry. The settings for demultiplexing were 0 mismatches in the barcode sequences.

### Bioinformatics

Bioinformatic analyses followed a published protocol (Busch et al. 2022). For computation of microbial core-diversity metrics, sequences were processed within the QIIME2 environment (v2024.10) (Bolyen et al. 2019). Amplicon Sequence Variants (ASVs) were generated from forward reads (truncated to 270 nt) with the DADA2 algorithm (Callahan et al. 2016). Contaminants were removed based on negative DNA extraction, PCR, and sequencing blanks with the {deconam} R package (Proctor et al. 2018). Representative ASVs were classified using the Silva v138.1 99% 16S rRNA gene database (Quast et al. 2013; Robeson et al. 2021) with the help of a primer-specific trained Naive Bayes taxonomic classifier. Mitochondrial, chloroplast and unassigned reads were removed, and singletons occurring only in one sample removed. Rarefaction curves based on species counts (Observed richness) and the Shannon diversity index were constructed for all samples (**Fig. S2**). The data were rarefied to the sample with the lowest count, a sampling depth of 4579 reads per sample. Alpha and beta diversity indices were calculated within QIIME2, further microbiome analyses and visualizations were done with MicrobiomeAnalyst (Dhariwal et al. 2017), {phyloseq} (McMurdie and Holmes 2013), {ampvis2} (Andersen et al. 2018), and {vegan} (Dixon 2003) R packages.

### Statistical analysis

All statistical analyses were performed in R (v4.3.3) (R Core Team 2024) and RStudio (v2023.12.1) (Posit team 2024) at a significance level of α = 0.05. Mean values reported as mean ± 1 SD. To assess differences in seawater parameters, and alpha diversity metrics we used Kruskal-Wallis tests, followed by pairwise Wilcoxon rank-sum post-hoc tests with Holm correction for multiple comparisons. Significance levels are indicated as: **** (p ≤ 0.0001), *** (p ≤ 0.001), ** (p ≤ 0.01), * (p ≤ 0.05), and ns (not significant, p > 0.05). Permutational multivariate analysis of variance (PERMANOVA) was conducted using the {vegan:adonis2} function. Significance of differences on boxplots were annotated directly with the {ggsignifi} package based on results of the Wilcoxon rank-sum test. Pairwise correlation coefficients were calculated for all average monthly environmental parameters using Spearman’s rank correlation. Correlation strength was classified based on the absolute value of the correlation coefficient: strong >0.7, moderate 0.7 ≥ R > 0.3, weak ≤0.3; and visualized with the package {ggcorrplot}.

## Results

### Repeated sponge sampling throughout seasons

In February 2022, around a wave breaker with natural populations of the sponge *H. panicea* in proximity to a beach at the northern German Baltic coast, 10 field stations were established at 1-3 m deep and with 1-2 m distance between them. Individual sponges were tagged for the long-term monitoring and monthly microbiome sampling (**Fig 1A**). In March 2022, four additional stations were established. Due to environmental conditions, stations established in February and March 2022 were lost and 12 new stations were established in April 2022. Stations on the outside of the wave breaker partially overlapped with the location of older stations. In August and September 2022, two new sponges were tagged and sampled at existing stations (gps15 and gps16). Based on the sampling success and continuity, stations were grouped into three categories: aborted - lost after March 2022 (10% sponge samples), partial - occasional failed sampling or loss of marked sponge individuals (28.5% sponge samples), and continuous - successful repeated sampling of marked sponges at recorded stations (61.5% sponge samples) (**Fig 1B**). Based on photo-documentation, there were observable differences in morphology throughout months and seasons at all stations of monitored sponges. Sponge biomass was seen to increase April to July 2022, suddenly decrease in August 2022 and continue to diminish until December 2022, after which in January 2023 an increase was observed again until May 2023 (**Fig 1C**). From the beginning of the monitoring in Feb 2022 until the end in May 2023, a total of 204 sponge and 57 seawater samples was obtained, with a median of 13 sponge and 4 seawater samples per month (**Table S1**).

**Figure 1.**
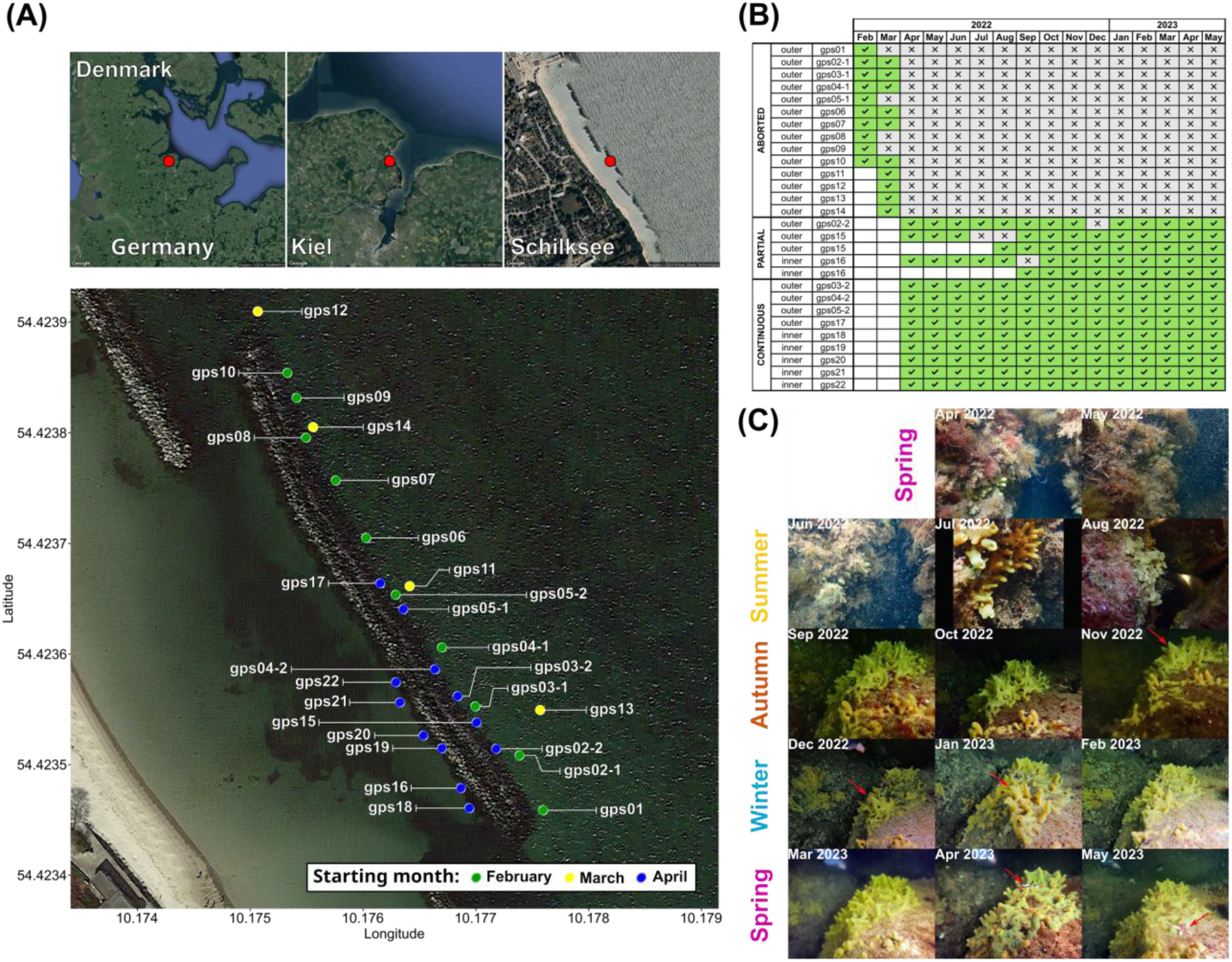
Monthly sampling design of tagged Halichondria panicea sponges. **(A)** Map of sampling location with individual field stations, established in February, March, and April 2022, where individual sponge colonies were repeatedly sampled. **(B)** Overview of sampling success at each field station over 16 months starting with Feb 2022 and ending in May 2023, with their location at the wavebreaker and sampling success. White = not established, green = successful sponge sampling, gray = lost station or failed sponge sampling. **(C)** Collage of monthly snapshots of a representative continuous field station (gps21) and the repeatedly sampled sponge individual from Apr 2022 to May 2023. Red arrows indicate drastic changes in the sponge colony and potential omnivory by sea stars.

Over the course of 16 months, we observed considerable seasonal and monthly variations in seawater temperature, salinity levels, light intensity, and daily light duration at the benthos throughout the year (**Fig. 2**). Annual seawater temperature minima occurred during winter, with the lowest average monthly temperature happening in February 2023 (4.59±0.28°C) and daily minimum in December (3.26°C), while maxima occurred in summer, with highest monthly averages (20.42±0.80°C) and the daily maximum (21.47°C) in August. The largest temperature fluctuations within a month (11.18-20.76°C) were recorded in summer (June), while winter (February 2022) showed the smallest fluctuations (4.90-5.31°C). Inconsistent with stable winter temperatures, the recorded minimum and large drop from average monthly values in seawater temperature in December (6.00±0.96°C) could indicate an upwelling or turbulent atmospheric event.

**Figure 2.**
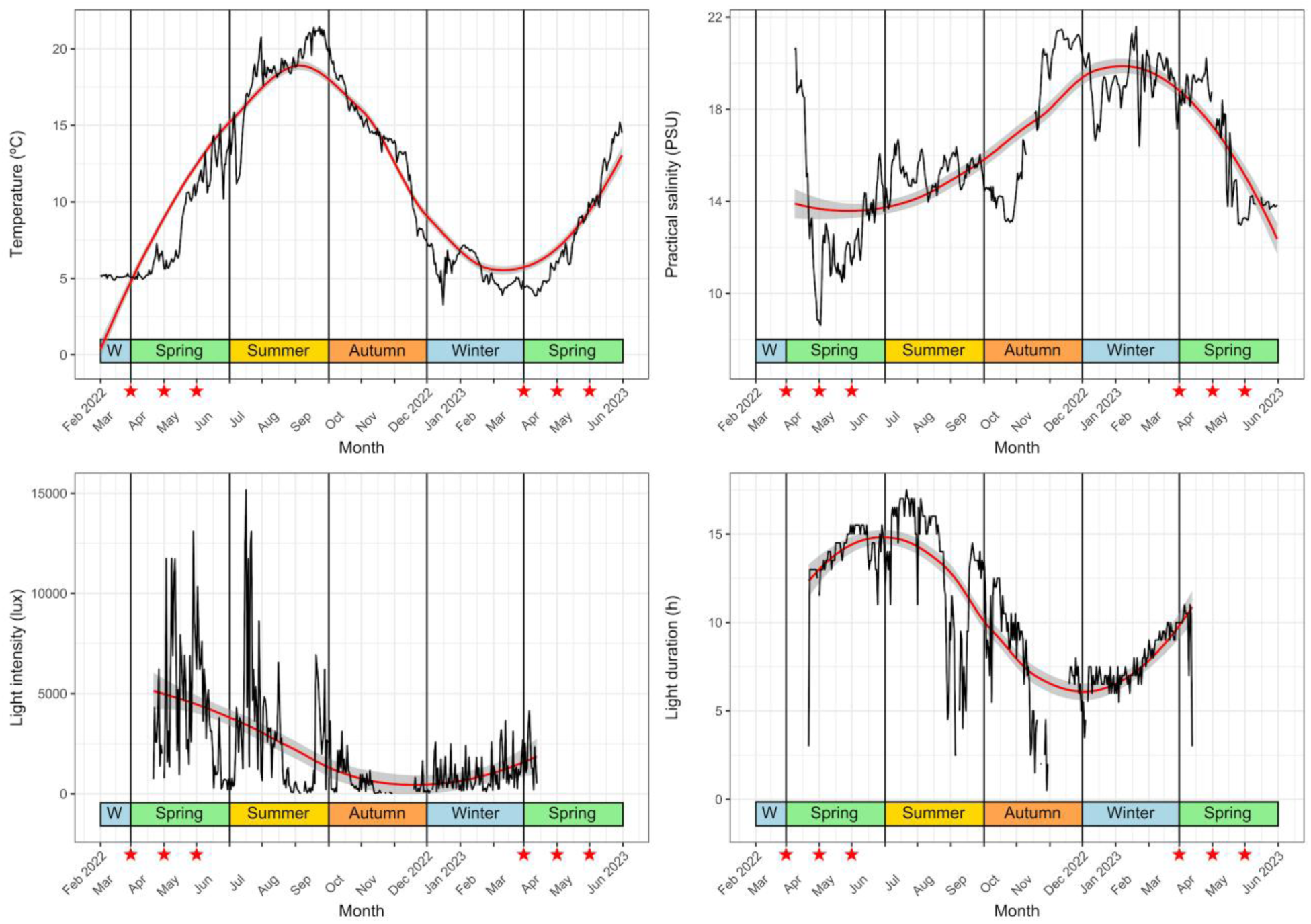
Annual trends in daily average seawater temperature, practical salinity, benthic light intensity, and total daily light duration from Feb 2022 to May 2023. Red line represents a LOESS (Locally Estimated Scatterplot Smoothing) fit, gray area as the 95% confidence interval. Red stars indicate months with sponge reproductive activity (Mar-May).

**Figure 3.**
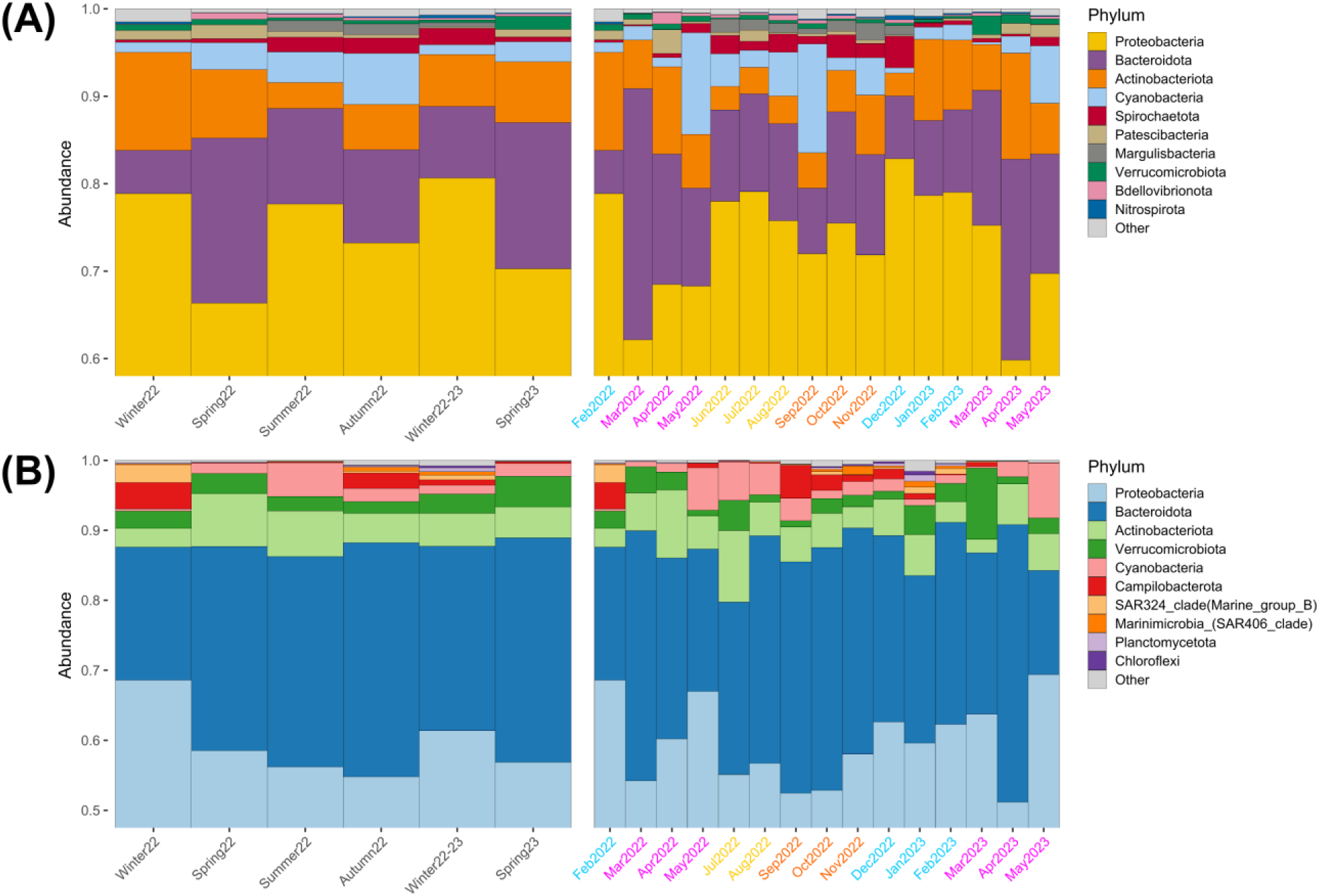
Taxonomic composition at the phylum level across seasons (Winter 2022 - Spring 23) and individual months (February 2022 - May 2023) over 16 months for sponge (A) and seawater (B) samples. Spring months (pink) coincide with reproductive timings.

**Figure 4.**
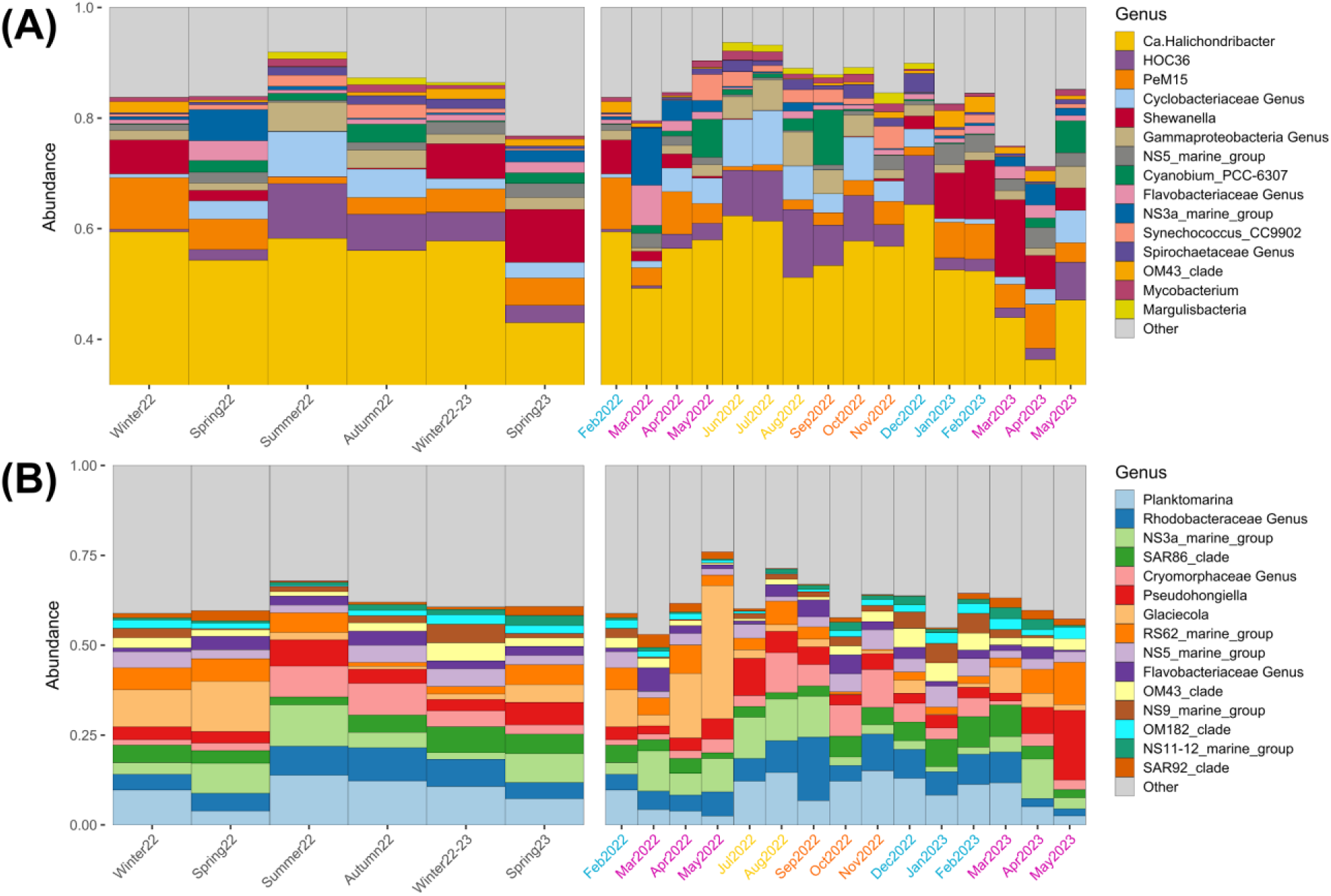
Taxonomic composition at the genus level across seasons (Winter 2022 - Spring 23) and individual months (February 2022 - May 2023) over 16 months for sponge (A) and seawater (B) samples. Spring months (pink) coincide with reproductive timings.

**Figure 5.**
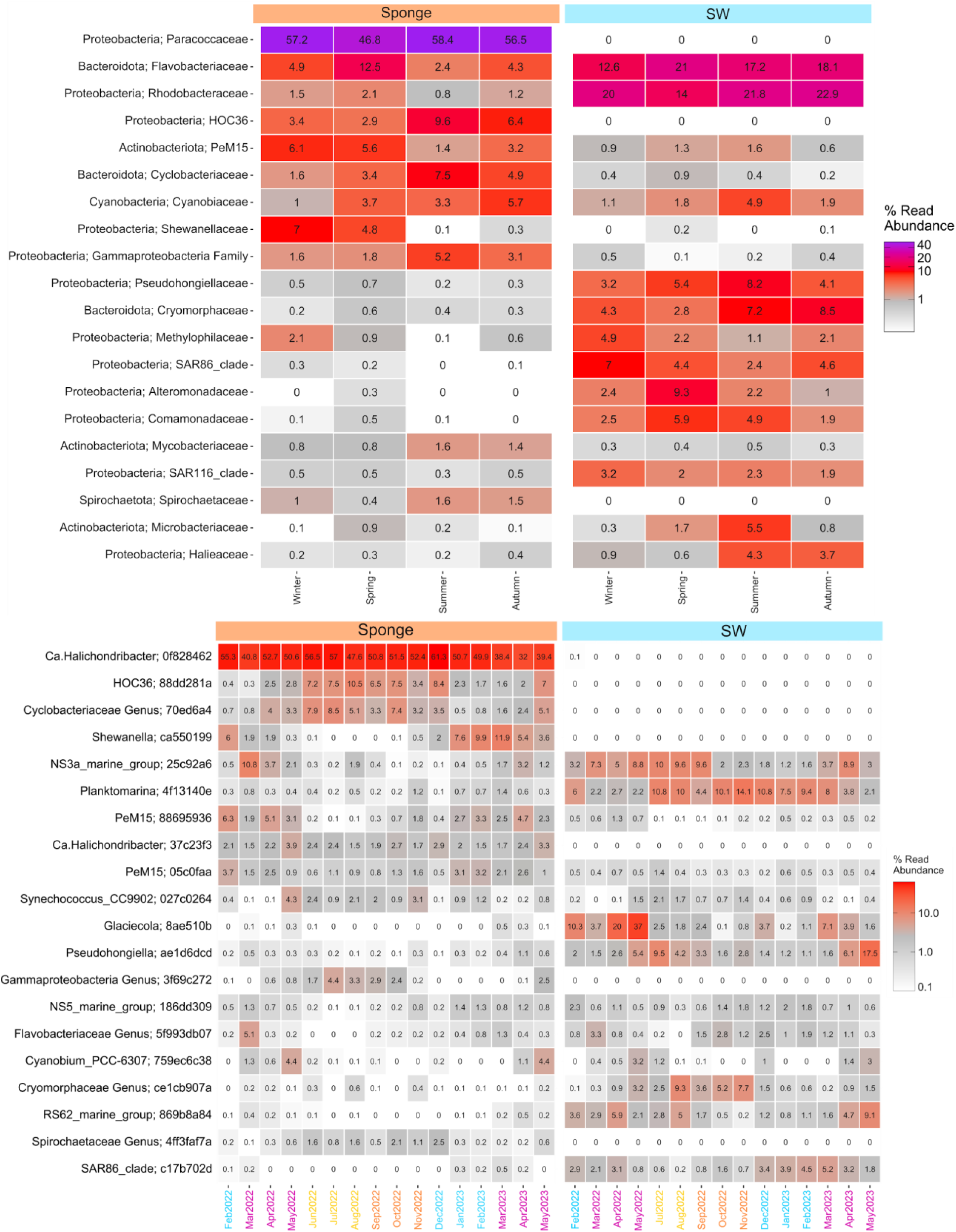
Heatmap of the bacterial taxonomic composition. Seasonal variation at the family level (top), and monthly variation at the ASV level (bottom) for sponge and seawater samples over a period of 16 months (Feb 2022 - May 2023). Individual months colored according to season. Spring months (pink) correspond to reproductive timings.

Salinity exhibited a different dynamic, with minima measured in spring for lowest average monthly data (11.39±1.08 PSU) and the daily maximum (8.62 PSU) recorded in April 2022, while maxima occurred in winter as the highest monthly average in November (20.93±0.478 PSU) and the daily maximum was recorded in January (21.60 PSU). Largest and smallest variations of salinity within a month also happened in spring, in March 2022 (8.85±20.66 PSU) and May 2023 (13.20±14.26 PSU) respectively. Increases in winter and decreases in spring-summer occur regularly in the Baltic, with sudden decreases between October and January likely related to freshwater discharge events and atmospheric disturbances.

Strong seasonal photoperiodicity was observed in relation to light measurements. Subsurface light intensity minima were recorded in autumn with lowest average monthly values in November (0.30±0.43 Klux) and daily minima in October and November (0.02 Klux), while maxima occurred in spring and summer with highest average monthly intensity recorded in April 2022 (5.83±3.59 Klux) and daily in June (15.17 Klux). Summer showed the largest variation within a month in June (0.22-15.17 Klux) and the smallest in autumn in October (0.02-1.17 Klux). Light duration followed a similar pattern, with minima in winter for average monthly data in November (5.60±2.65 h) and daily minima again in October and November (0.5 h), while maxima for the monthly average (15.72±1.65 h) and daily maximum (17.5 h) were in June. Largest fluctuations occurred in summer for July (4.5-17.0 h) and the smallest in winter for February 2023 (8.0-10.0 h). Slight deviations of subsurface from atmospheric photoperiod values and timings might have occurred, due to biofouling and the sensitivity of the logger’s light sensor during regular cleaning procedures.

Analyzing the monthly averages of environmental seawater parameters (**Fig S1**) reveals distinct seasonal and monthly variations that are characteristic of temperate marine environments in the Baltic. Winter (February) stands out with high nutrient levels (nitrite, nitrate, phosphate, and silicate) and the lowest temperature. Spring (April) is characterized by peak light intensity and oxygen levels but minimal salinity. Throughout summer, the longest sunlight duration, lowest nutrient levels and oxygen saturation were observed, with peak temperatures. Autumn (November) marked high salinity but minimal light intensity and sunlight duration, while winter again (December) had the lowest pH. There were no significant differences between the spring seasons in different years, except for salinity (p < 0.001), light intensity (p < 0.05), and light duration (p < 0.001), but overall significant differences between the colder (autumn-winter) and warmer (spring-summer) months were detected in all parameters with daily measurements (**Table S2**).

Our correlation analysis between annual patterns of environmental parameters in seawater (**Fig S2)**, showed strong correlations among monthly averages in several variables. Temperature, salinity, light intensity, and nitrite were identified as key parameters for further analysis, as they collectively capture the primary physical, chemical, and biological processes in the dataset. Temperature strongly correlated with pH (R = 0.747), dissolved oxygen (R = -0.794), and nitrate (R = -0.821). Salinity showed strong negative correlations with light intensity (R = -0.794) and light duration (R = -0.824). Nitrite strongly correlated with silicate (R = 0.868) and nitrate (R = 0.791), while its negative correlation with light duration (R = -0.803) reflects phototrophic nutrient uptake. Light intensity showed a strong positive correlation with light duration (R = 0.921) and strong negative correlations with phosphate (R = -0.909) and silicate (R = -0.724). Generally, the dissolved nutrient measurements showed strong interdependencies.

### Seasonal Community Shifts

After sequencing, 200 sponge and 53 seawater samples produced usable data. In order to keep as many samples as possible to preserve paired information for a repeated measures design, at least 4500 reads were chosen as the minimum number of sequences needed to pass rarefaction criteria, which resulted in only one discarded sponge sample (**Fig. S3**). A total of 2833 ASVs was recovered from 16 months of sponge and seawater sampling. Based on the observed richness (species/ASV count), sponge samples (384.53±159.73) were on average slightly richer than seawater samples (339.85±89.47), and had more extreme values with up to two-fold higher richness than seawater samples (**Fig. S4**). A trend that was observed throughout most monthly collections, with the lowest number of observed species for sponge detected in June (226.92±92.05) and seawater samples in July (186.00±17.05), while the highest was in January (533.36±169.56; 480.50±7.72) respectively. Sponge samples also consistently showed greater variability in richness as seen with a higher mean interquartile range (168.38±73.83) than seawater samples (23.23±20.40) throughout the months.

The microbiome of *H. panicea* exhibited distinct seasonal profiles. During summer, the community was dominated by symbionts favoring higher temperatures and increased primary productivity. Winter communities, in contrast, were composed of bacteria adapted to lower temperatures and reduced metabolic activity. These seasonal shifts were characterized by rapid transitions between summer and winter states during spring and fall. For instance, the summer-dominant bacteria included members of the classes Alphaproteobacteria and Gammaproteobacteria, which are known for their metabolic versatility and ability to thrive in nutrient-rich conditions. In winter, the community was dominated by Bacteroidetes and Planctomycetes, which are more adapted to colder, nutrient-poor environments.

These findings suggest that the seasonal shifts in the *H. panicea* microbiome are driven by changes in environmental conditions, in particular, temperature and nutrient availability. The rapid transitions between summer and winter states indicate that the bacterial community is highly responsive to changes in its environment, with specific taxa showing preferences for different seasonal conditions. This responsiveness highlights the dynamic nature of sponge-associated microbial communities and their potential to adapt to changing environmental conditions, where closely related bacterial strains showed seasonal preferences, emphasizing the adaptability of marine bacterial communities to environmental fluctuations.

### Alpha diversity

The PERMANOVA results (**Fig 6**, **Table S3**) for Shannon diversity indicate that both sample type and season strongly influence the variation in diversity. Sample type accounts for the largest proportion of variance, explaining 65.31%, which is highly significant and season explains a smaller but still significant portion of the variance (8.93%). The interaction between season and sample type also contributes significantly, though its effect size is relatively small at 3.65%, indicating that the influence of sample type varies depending on the season. Shannon diversity for SW samples was higher than in sponges during all seasons. For ACE, a richness estimator, season plays a dominant role, explaining 32.88% of the variance, whereas the sample type and interaction contributed negligibly (0.90% and 0.025% respectively). These results suggest that richness is primarily influenced by seasonal changes and is negligible between sample types. Faith’s PD, a phylogenetic diversity metric, shows that season is again a major driver, explaining 36.13% of the variance with statistical significance. While sample type didn’t have a significant effect, the interaction between season and sample type is significant, though contributing only 2.44%. This suggests that phylogenetic diversity is influenced by season and that there may be subtle differences in sample types during specific seasons. While no differences were detected in Autumn and Winter, diversity was significantly lower in SW compared to sponges in spring and summer. Comparing the metrics, Shannon diversity highlights the dominant effect of sample type, whereas ACE and Faith’s PD emphasize seasonal effects. Richness (ACE) and phylogenetic diversity (Faith’s PD) are minimally influenced by sample type, in contrast to Shannon diversity, which captures both richness and evenness and shows a strong sample type effect. The distinction between the two suggests that differences in evenness, such as variations in the distribution of abundances among taxa, are significant drivers of the patterns observed in Shannon diversity but are not reflected in ACE.

**Figure 6.**
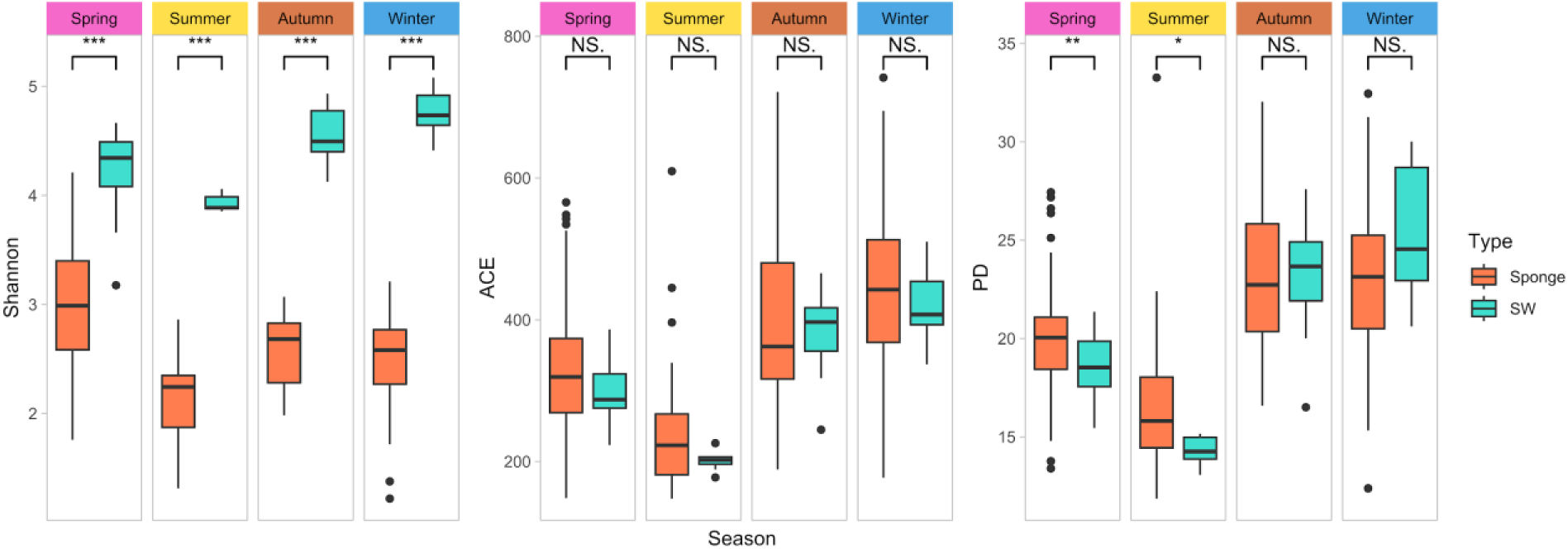
Seasonal alpha diversity measures (Shannon, ACE, Faith’s PD index) for sponge and seawater samples over 16 months (Feb 2022 - May 2023). Significant differences detected with Wilcoxon rank sum test as * <0.05, ** <0.01, *** <0.001.

**Figure 7.**
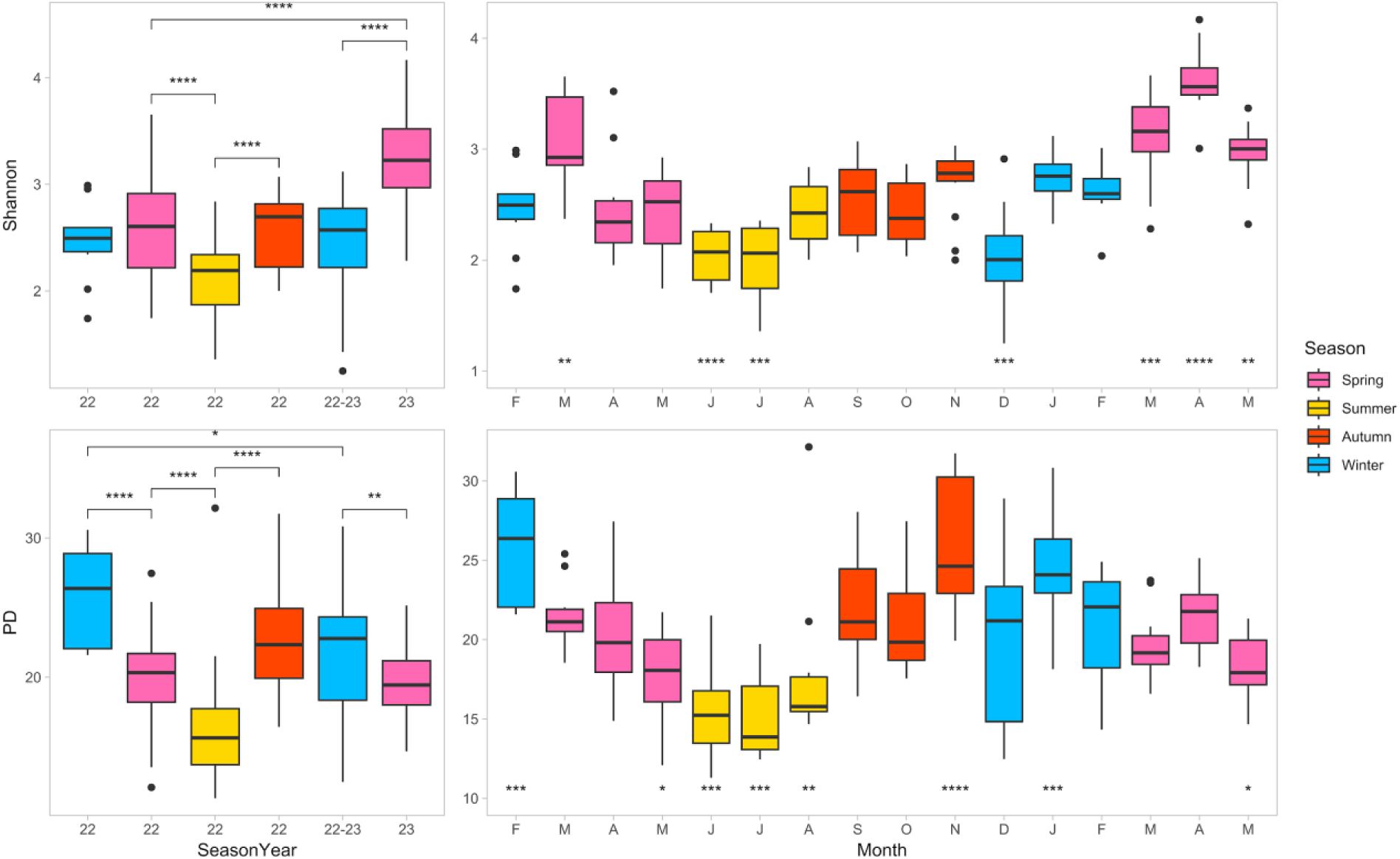
Alpha diversity measures for individual seasons (Winter 2022 - Spring 2023) and months (Feb 2022 - May 2023) for sponge samples expressed with Shannon’s (top) and Faith’s PD index (bottom). Colors correspond to seasons.

**Figure 8.**
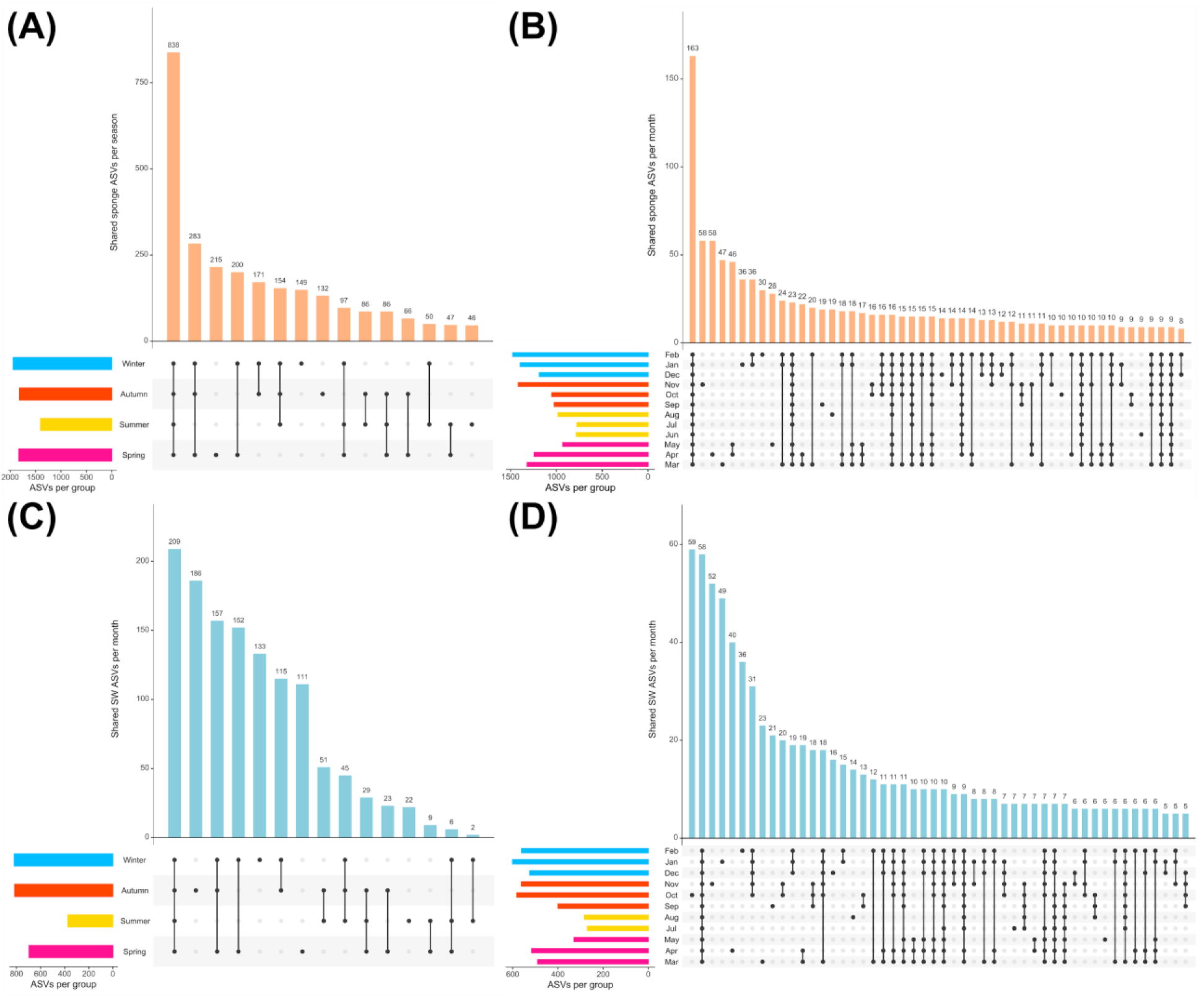
Seasonal and monthly sharing of ASVs. Seasonal (A) and monthly (B) for sponge samples, and seasonal (C) and monthly (D) grouping for seawater samples. Months colored according to seasons.

**Figure 9.**
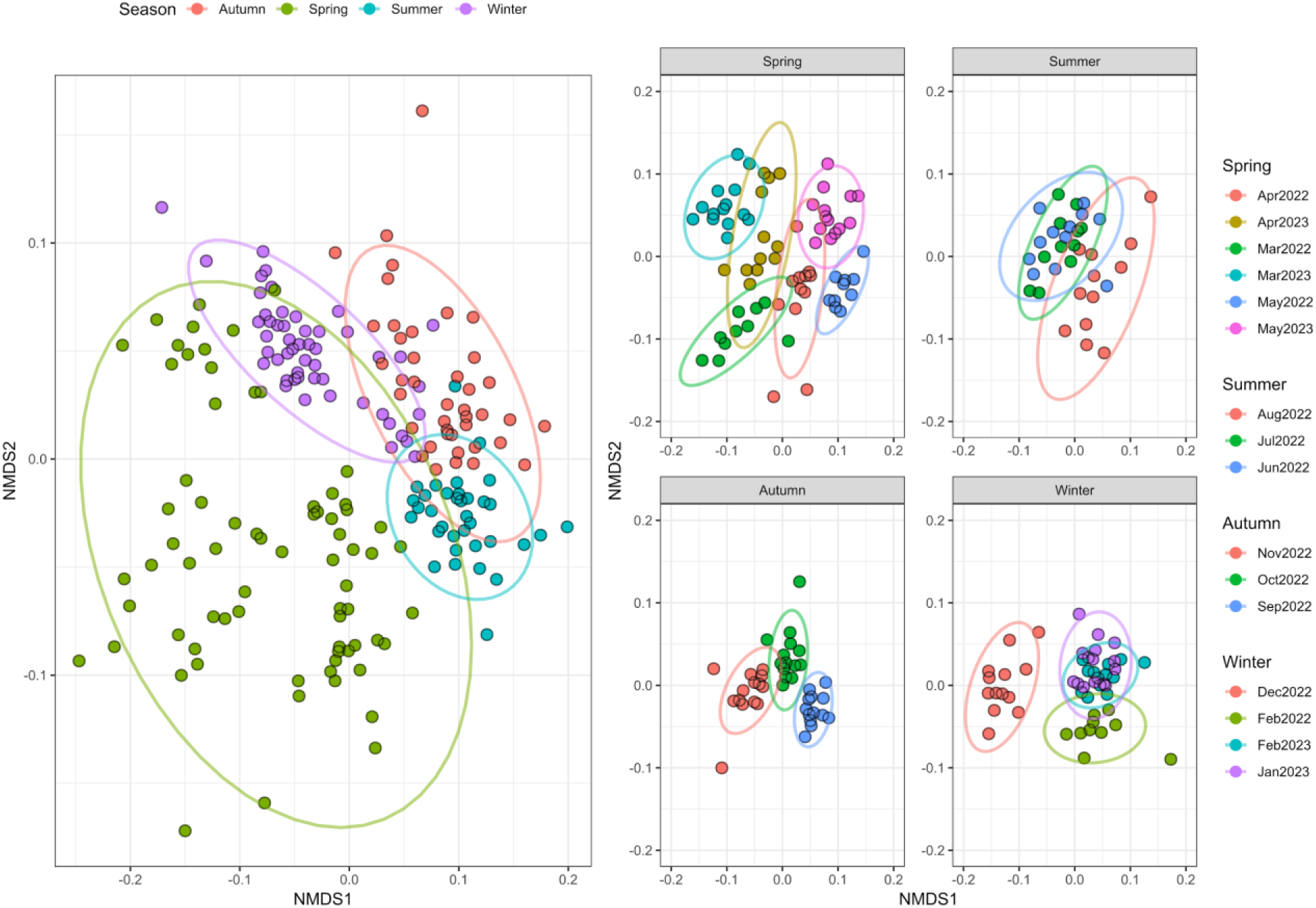
Separation of seasons and months according to seasons according to an NMDS analysis for sponge samples.

The repeated increases in Shannon, ACE, and Faith’s PD indices during November and January suggest a seasonal or environmental factor positively influencing microbial diversity during these months, particularly in sponge samples. This could be related to changes in water temperature, nutrient availability, or other seasonal events that promote microbial growth and diversity. The sharp decline observed in December across these indices could indicate a sudden change in environmental conditions, such as temperature drops or nutrient depletion, which adversely affects microbial richness and diversity. April consistently shows the lowest diversity values, which suggests a recurring stressor such as a cyclic event that negatively impacts the bacterial community, such as changes in environment chemistry or host physiology. The consistent dips observed across all alpha diversity indices (Fisher, Shannon, Simpson, ACE, and Chao1) during March, April, May 2022, and December 2022 suggest that the microbial communities in both seawater and sponge samples are resilient but periodically undergo stress (**Fig S5**). This pattern appears to affect all three indices similarly, indicating that the factors driving these changes impact overall richness and diversity rather than just specific aspects of the bacterial community.

The decline outside of the winter months suggests a loss of species (richness), particularly rare or less abundant ASVs. The stability in Shannon and Simpson indices implies that the overall structure of the community, including the relative abundances of the remaining species, is not changing drastically. Even though some species are lost, the remaining species are evenly distributed. Common or dominant species are resilient to these changes, maintaining the overall balance and evenness of the community. The consistent dips observed during March, April, May 2022, and December 2022 suggest that the microbial communities in both seawater and sponge samples are resilient but periodically undergo stress. This resilience might be crucial for the long-term stability of the sponge-microbiome association, allowing the microbiome to bounce back after unfavorable conditions.

### Beta diversity

The assessment of community differences using Bray-Curtis and weighted Unifrac distances between microbiome samples from sponges and seawater, we observed monthly and intersample variability with distinct patterns (**Fig 10**). The lowest average Unifrac distance within seawater samples was observed in February 2023 (0.03±0.004), indicating a high similarity among the microbial communities in these samples. In contrast, March 2022 exhibited the highest variability (0.23±0.08), suggesting a significant temporal divergence in the seawater microbiome. This month also recorded the greatest variability in both seawater-seawater and seawater-sponge comparisons, indicating substantial fluctuations in bacterial community compositions, possibly driven by seasonal environmental changes. The comparisons between seawater and sponge microbiomes showed the lowest similarity in October 2022, with the highest recorded average Unifrac distance of 0.43±0.02. This suggests a large distinction between the microbial communities of sponges and their surrounding seawater during this period. Conversely, the closest interaction occurred in March 2023 (0.28±0.05), possibly influenced by shared ecological or environmental conditions contributing to higher similarity.

**Figure 10.**
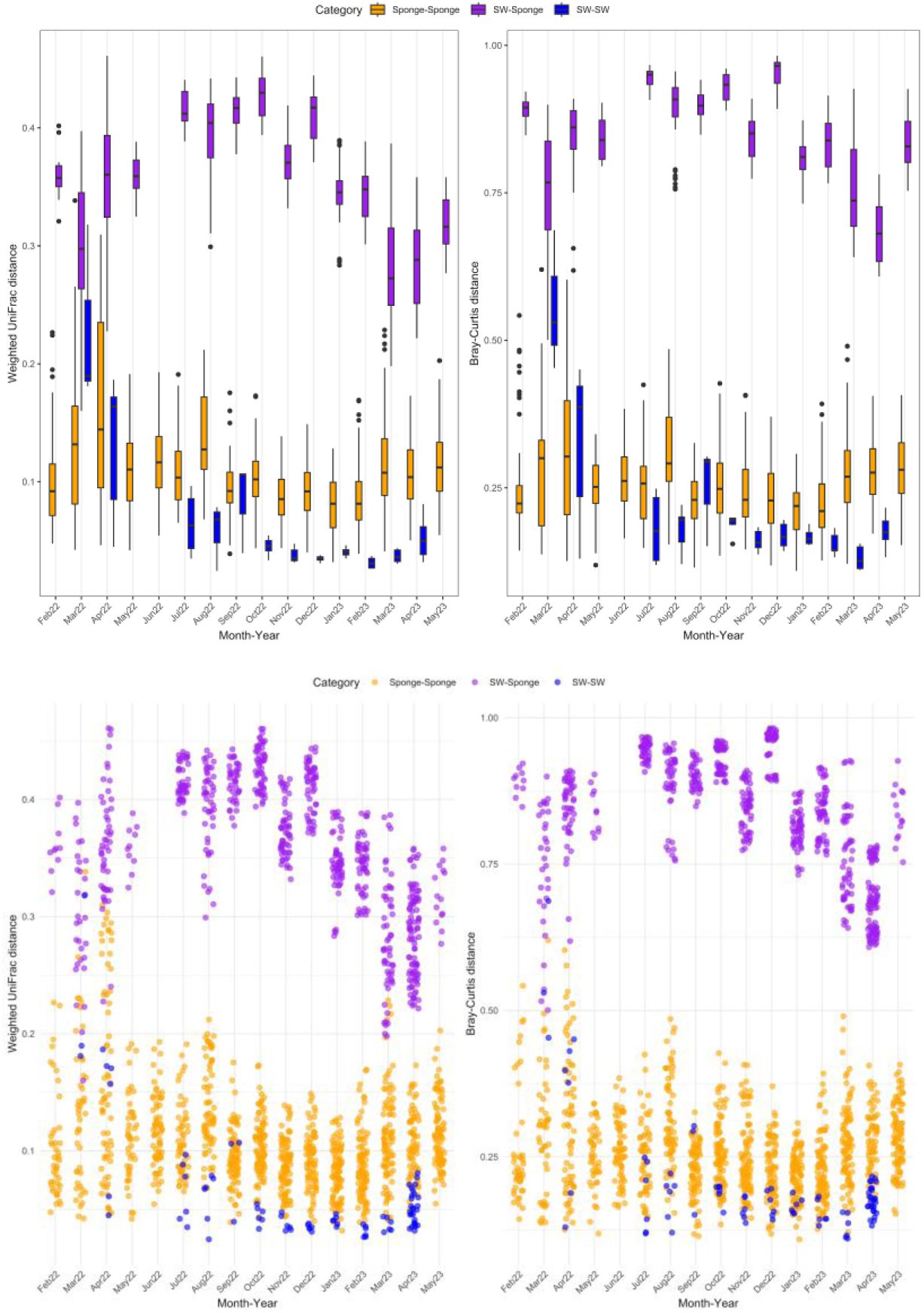
Bacterial community dissimilarity between sponges and seawater (SW) within 16 months. Box/jitter-plot of pairwise weighted Unifrac (left) and Bray-Curtis distances (right) calculated between sponge-sponge, sponge-seawater, and seawater-seawater samples for each month.

**Figure 11.**
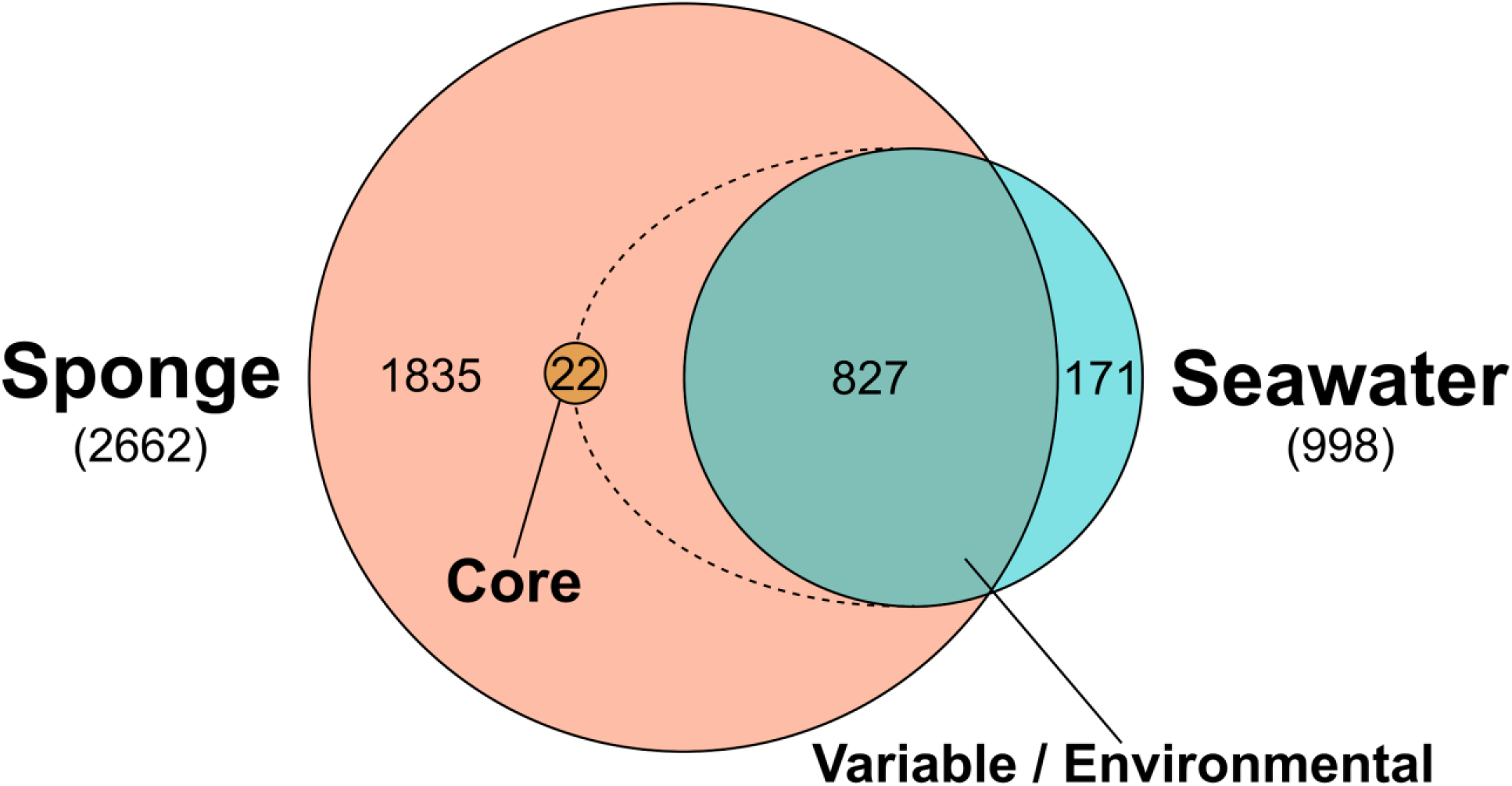
The stratification of sponge and seawater ASVs in sponge- and seawater-specific groups, variable/environmental ASVs contributed by seawater to the sponge microbiome, and the core sponge microbiome.

Among sponge samples, January 2023 displayed the most similar microbial profiles (0.08±0.02), indicating a stable sponge microbiome during this month. April 2022, however, showed the greatest diversity (0.16±0.08), aligning with the highest spread in bacterial community structure observed among all sponge samples. This period may reflect ecological disturbances or significant biotic shifts impacting sponge-associated microbial communities.

### Symbiont dynamics

*Candidatus* Halichondribacter symbioticus remained constant throughout the year, but its relative abundance fluctuated with the seasons. This symbiont peaked in abundance during the summer and winter months, when environmental conditions favored higher metabolic activity and nutrient availability. In contrast, its abundance decreased slightly during spring, reflecting the higher metabolic demands of the sponge host during this period which coincides with reproductive development. The seasonal fluctuations in the abundance of *Ca.* Halichondribacter symbioticus suggest that its interactions with other microbial members are not modulated by environmental conditions.

### Seasonal dynamics of individual taxa and environmental drivers of diversity

Examining the seasonal dynamics of individual taxa revealed that certain bacterial groups exhibited clear seasonal preferences. For example, members of the genus HOC36 were predominantly found during the summer months, while Shewanella species were more abundant in winter. The abundance of HOC36 in summer, for instance, could be linked to its ability to utilize the increased dissolved organic matter from primary production, while Shewanella might be more suited to the lower temperatures and nutrient conditions of winter. Environmental drivers such as temperature, salinity, pH, and light intensity were closely linked to the observed patterns of microbial diversity. Correlation analyses indicated that temperature was the primary driver of seasonal changes in community composition, with significant positive correlations between temperature and the abundance of summer-associated taxa. Conversely, nutrient availability, particularly nitrate and phosphate concentrations, showed strong correlations with the abundance of winter-associated taxa. These findings suggest the importance of environmental variables in shaping bacterial community dynamics and highlight the intricate relationships between environmental conditions and microbial diversity. The seasonal dynamics of individual taxa were further supported by the observation of rapid transitions between community states during spring and fall. These transition periods were characterized by a mix of summer and winter taxa, suggesting that microbial communities are highly responsive to changing environmental conditions. The rapid shifts in community composition during these periods emphasize the dynamic and adaptive nature of sponge-associated microbial communities. Similar rapid transitions have been documented in other marine microbial studies, indicating a common ecological strategy among marine bacteria to quickly respond to environmental changes.

### Core microbiomes

## Discussion

The seasonal dynamics of the *H. panicea* microbiome observed in this study provide valuable insights into the ecological and functional roles of sponge-associated microbial communities. Our findings reveal that the microbiome of *H. panicea* undergoes significant shifts in community composition and functional gene expression in response to seasonal environmental changes. These results are consistent with previous studies that have documented similar patterns in marine bacterial communities, which demonstrated seasonal switching between sponge-specific marine bacteria.

One of the key findings of this study is the distinct separation between summer and winter microbial communities within the *H. panicea* sponge. Summer communities were characterized by higher abundances of bacteria associated with nutrient cycling and primary production, while winter communities showed increased abundances of bacteria involved in stress response and nutrient scavenging. This seasonal partitioning suggests that specific bacterial taxa are adapted to the environmental conditions prevalent during particular seasons, supporting the idea of niche specialization among sponge-associated microbes. The rapid transitions between summer and winter microbial communities observed during spring and fall highlight the dynamic nature of the *H. panicea* microbiome. These transition periods were characterized by a mix of summer and winter taxa, indicating that the bacterial community is highly responsive to changing environmental conditions. Similar rapid transitions have been documented in other marine microbial studies, suggesting a common ecological strategy among marine bacteria to quickly respond to environmental changes. This adaptability is likely a key factor in the resilience of the sponge microbiome to environmental fluctuations. The dominance of *Candidatus* Halichondribacter symbioticus throughout the year confirms its importance in the sponge-microbe symbiotic relationship. However, the fluctuating abundance of this symbiont suggests that it interacts dynamically with other bacterial community members, modulating its role in response to environmental conditions. Understanding the dynamics of key symbionts like *Candidatus* Halichondribacter symbioticus is crucial for predicting the impacts of environmental changes on sponge health and function.

Seasonal variations in sponge microbial community composition have been observed across regions such as the Mediterranean Sea, Caribbean Sea, New Zealand, and Red Sea. Consistent with these findings, this study revealed seasonal fluctuations in sponge-associated bacterial composition and diversity. In *H. panicea*, bacterial diversity and composition differed significantly across seasons. In October, sponges exhibited the highest bacterial abundance but the lowest diversity, whereas July samples showed the highest diversity and lowest relative abundance. These patterns correlated with temperature changes, as higher temperatures in July were linked to increased Cyanobacteria and *Bacteroidota* abundance. Cyanobacterial growth in summer was driven by elevated sunlight needed for photosynthesis, while the abundance of symbiotic *Bacteroidota* correlated with higher temperatures and nutrient availability. The seasonal dynamics of individual taxa and the environmental drivers of diversity observed in this study underscore the importance of considering temporal variability in microbial ecology. Our findings demonstrate that temperature and nutrient availability are key drivers of bacterial community composition and function, highlighting the intricate relationships between environmental conditions and microbial diversity. These results have important implications for understanding the resilience and adaptability of marine sponges and their associated microbiomes in the face of environmental change. Overall, our results demonstrate that the *H. panicea* microbiome is highly dynamic and strongly influenced by seasonal variations in environmental conditions. The clear seasonal patterns observed in both community composition and dominance of specific bacterial groups highlight the adaptive capacity of the bacterial community to changing environmental conditions. These dynamics are crucial for predicting how marine sponges and their associated microbiomes will respond to future environmental changes, particularly in the context of global climate change.

Future research should focus on the long-term baseline monitoring of sponge-associated microbial communities to gain a more comprehensive understanding of their temporal dynamics. Additionally, exploring the impacts of other environmental drivers, such as pH and salinity, on bacterial community composition and function will provide further insights into the factors influencing microbial diversity. Investigating the mechanisms underlying the interactions between key symbionts and other community members will also be crucial for understanding the complexity of sponge-microbe symbiotic relationships. In conclusion, this study provides a detailed analysis of the seasonal dynamics of the microbiome associated with the marine sponge *Halichondria panicea*. Our findings reveal distinct seasonal patterns in bacterial community composition and function, driven by environmental factors such as temperature and nutrient availability. The rapid transitions between summer and winter communities during spring and fall highlight the dynamic nature of sponge-associated microbial communities and their ability to adapt to changing environmental conditions. Understanding these dynamics is crucial for predicting the resilience and adaptability of marine sponges and their associated microbiomes in the face of ongoing climate change.

## Conclusion

Our study contributes to this understanding by providing detailed insights into the temporal patterns of the *H. panicea* microbiome and its functional responses to seasonal variations. Future research should focus on the long-term monitoring of these communities and the exploration of other environmental drivers, such as pH and salinity, to gain a more comprehensive understanding of the factors influencing bacterial community dynamics. The seasonal switching between sponge-specific marine bacteria within the *H. panicea* sponge microbiome highlights the intricate relationship between environmental conditions and bacterial community structure and function. This study displays the importance of considering temporal dynamics in microbial ecology and provides a foundation for future research on the impacts of environmental change on marine symbiotic relationships. Understanding these dynamics is essential for predicting the resilience and adaptability of marine ecosystems in the face of ongoing climate change.

## Acknowledgements

We acknowledge Markus Zimmerer and the CAU Forschungstauchzentrum for sponge collection and the Competence Center for Genomic Analysis (CCGA) Kiel for amplicon sequencing. We thank members of the CRC1182 “Origin and Function of Metaorganisms” and members of the Hentschel lab for useful discussions. This project is supported by funding from the DFG (“Origin and Function of Metaorganisms”, CRC1182-TP C04).

