## Supplemental Figures S1-S10 for "Annual community patterns in the *Halichondria panicea* sponge microbiome are characterized by seasonal switching between sponge-specific marine bacteria"

### Figure S1: Monthly environmental measurements

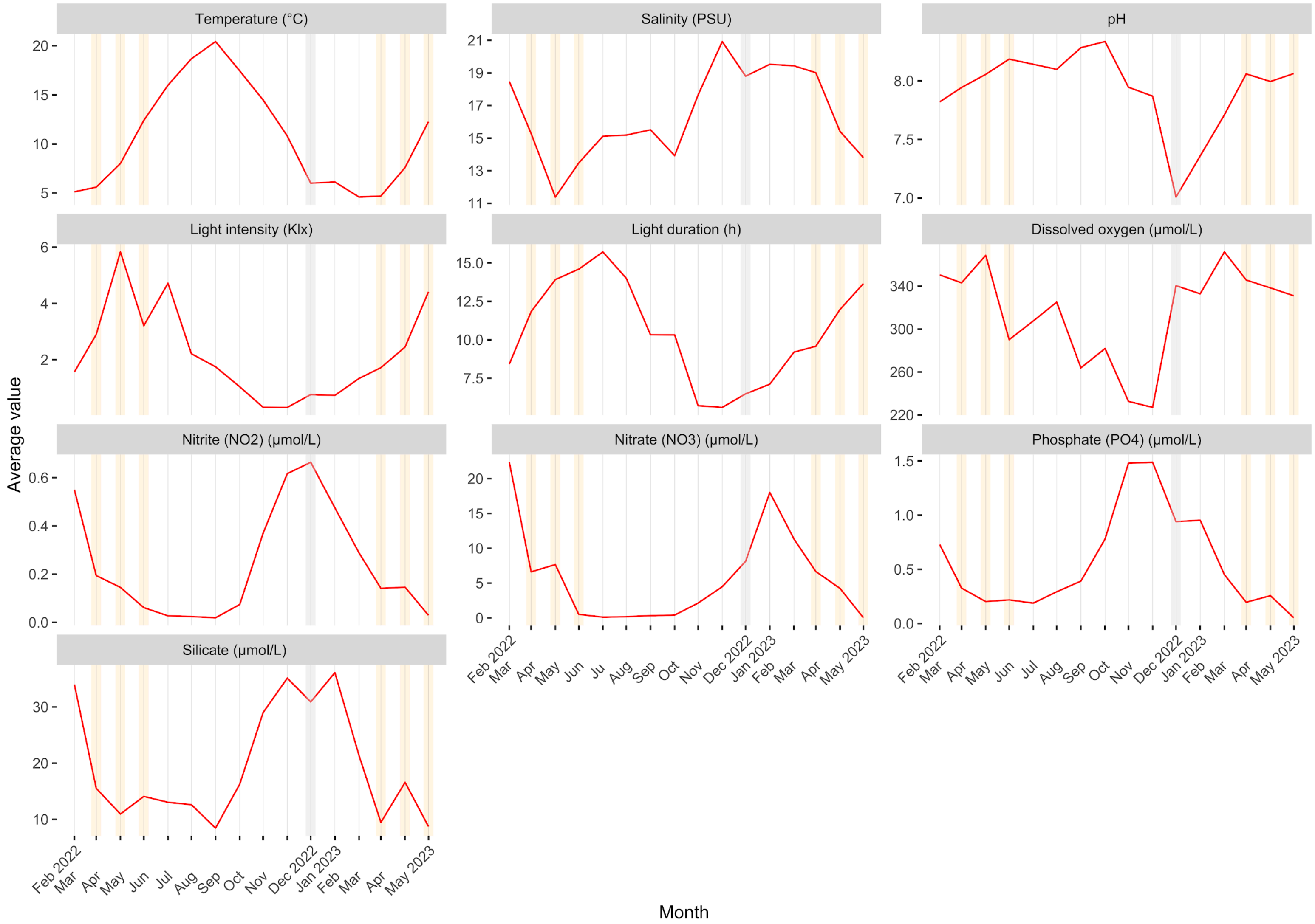


**Figure S1.** Annual dynamics in marine abiotic and biotic environmental parameters from Feb 2022 to May 2023. Averaged monthly measurements over 16 months relating to physical and biological parameters of seawater - temperature, light intensity, light duration, salinity, pH, dissolved oxygen, and nutrients ($NO_2$, $NO_3$, $PO_4$, silicate). Gray bar indicates Dec 2022, orange bars indicate months with sponge reproductive activity (Mar-May).

### Figure S2: Monthly environmental correlations

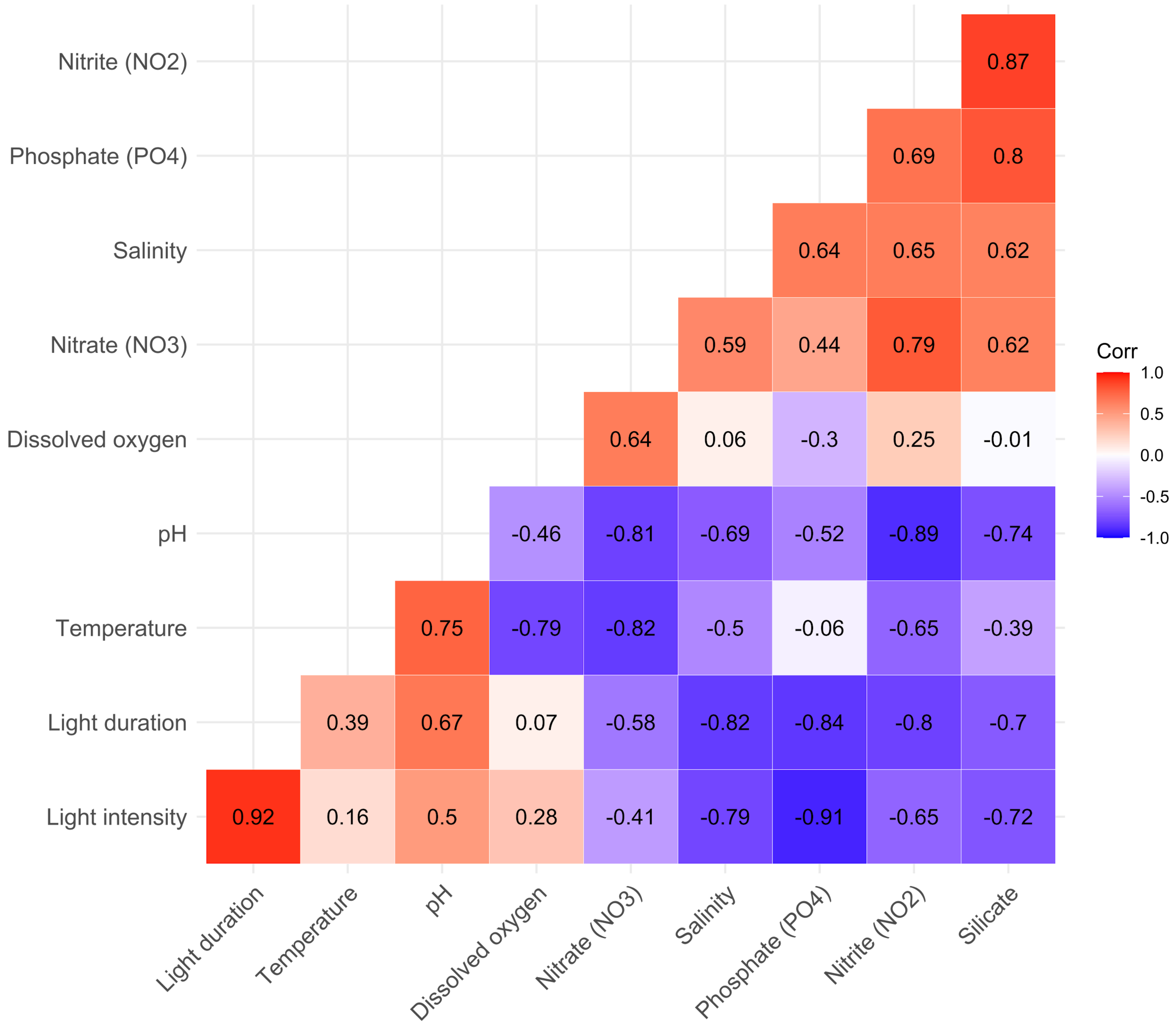


**Figure S2.** Correlation matrix of pairwise Spearman's rank correlation coefficients among annual marine environmental parameters of seawater based on monthly averages between Feb 2022 and May 2023. Colors indicate the strength and direction of the correlation: warmer shades (red) for positive correlations, cooler shades (blue) for negative correlations, and white for no correlation.

### Figure S3: Rarefaction curve

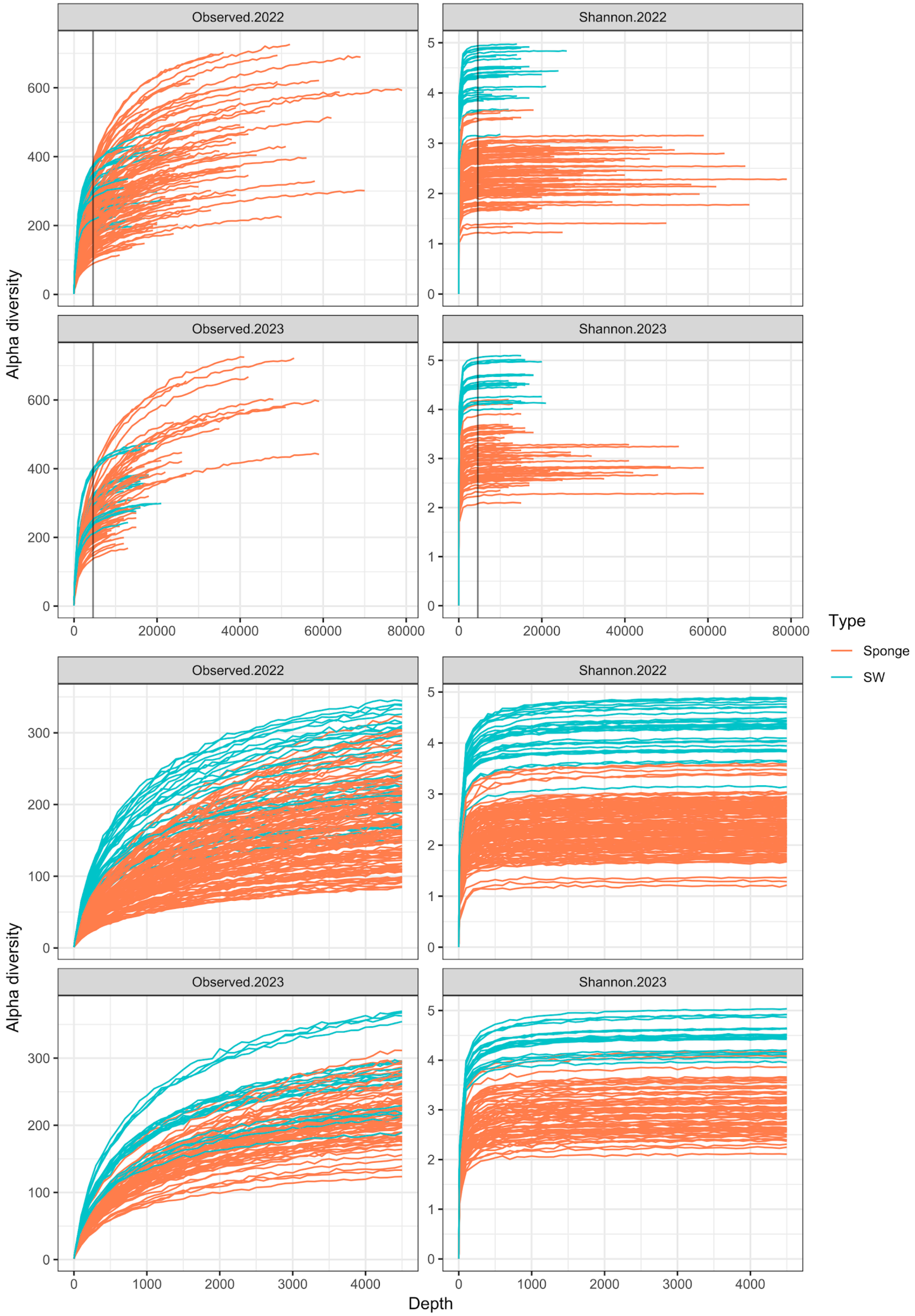


**Figure S3.** Amplicon sequence variation (ASV) richness rarefaction curves based on species counts (Observed) and Shannon diversity index (Shannon). Each line shows a sponge sample within the treatment category with error bars corresponding to the standard deviation at subsampled depth intervals. (Upper) Rarefaction curves before subsampling with the lowest sample depth indicated with a black vertical line (4500 reads). (Lower) Rarefaction curves after subsampling at 4500 reads.

### Figure S4: Sample richness

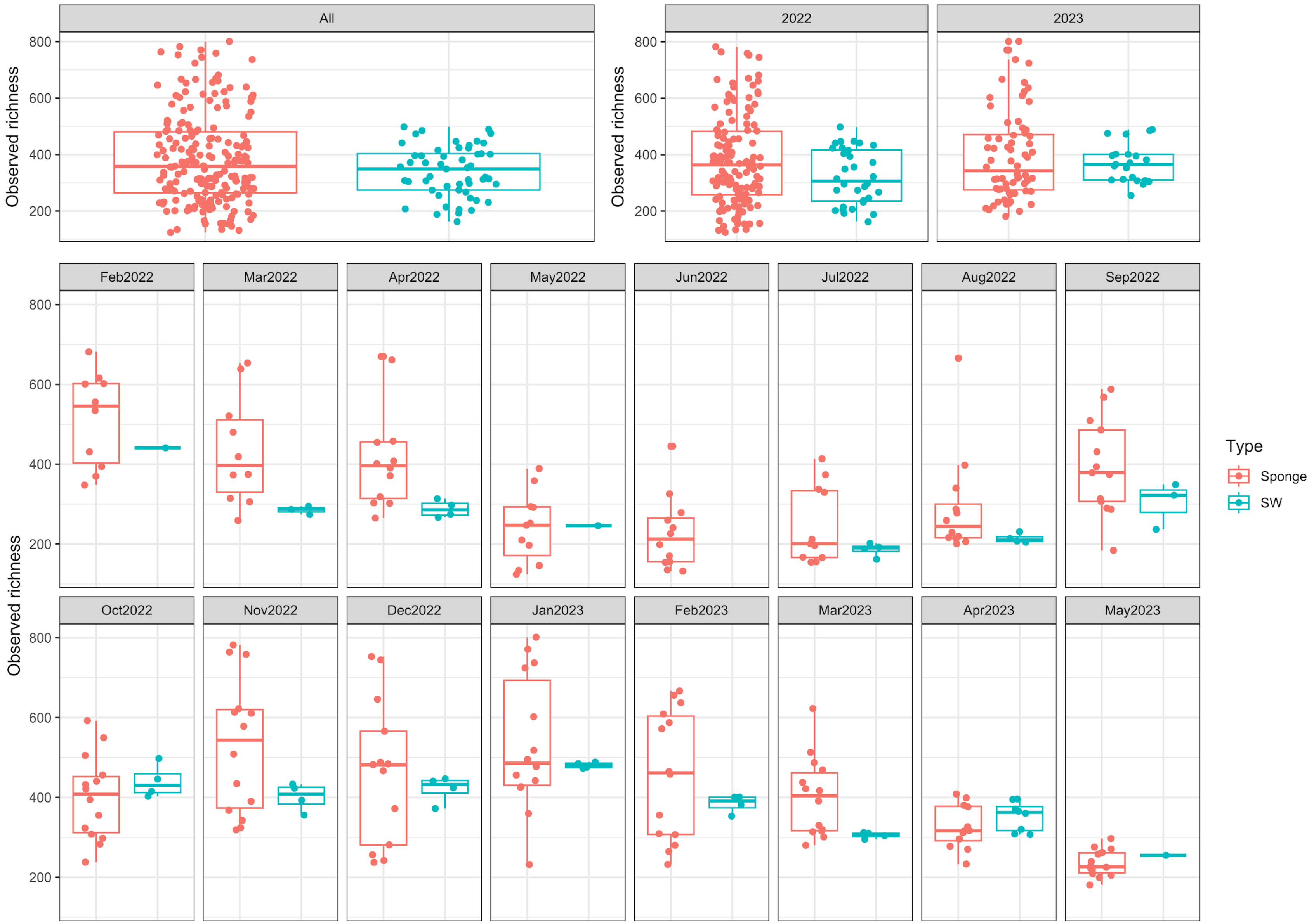


**Figure S4.** Sample richness distributions for all sponge and seawater samples, stratified by sampling year and months. Boxplots show median, interquartile ranges, minimum and maximum values for each group.

### Figure S5: Taxonomic composition for sponge and SW

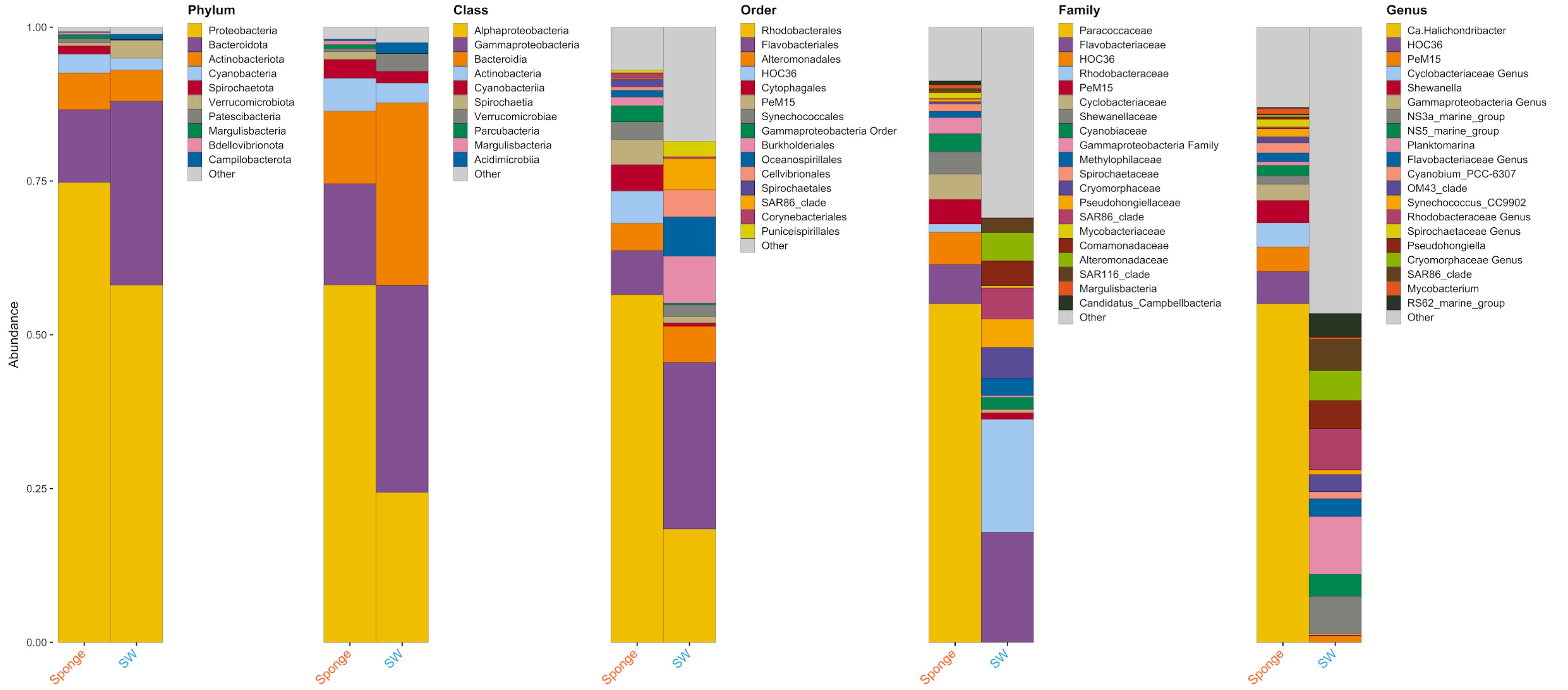


**Figure S5.** Taxonomic composition of sponge and seawater samples at different taxonomic levels (phylum-genus), for the top 10 (phylum-class), 15 (order), and 20 (family-genus) taxa.

### Figure S6: Alpha diversity

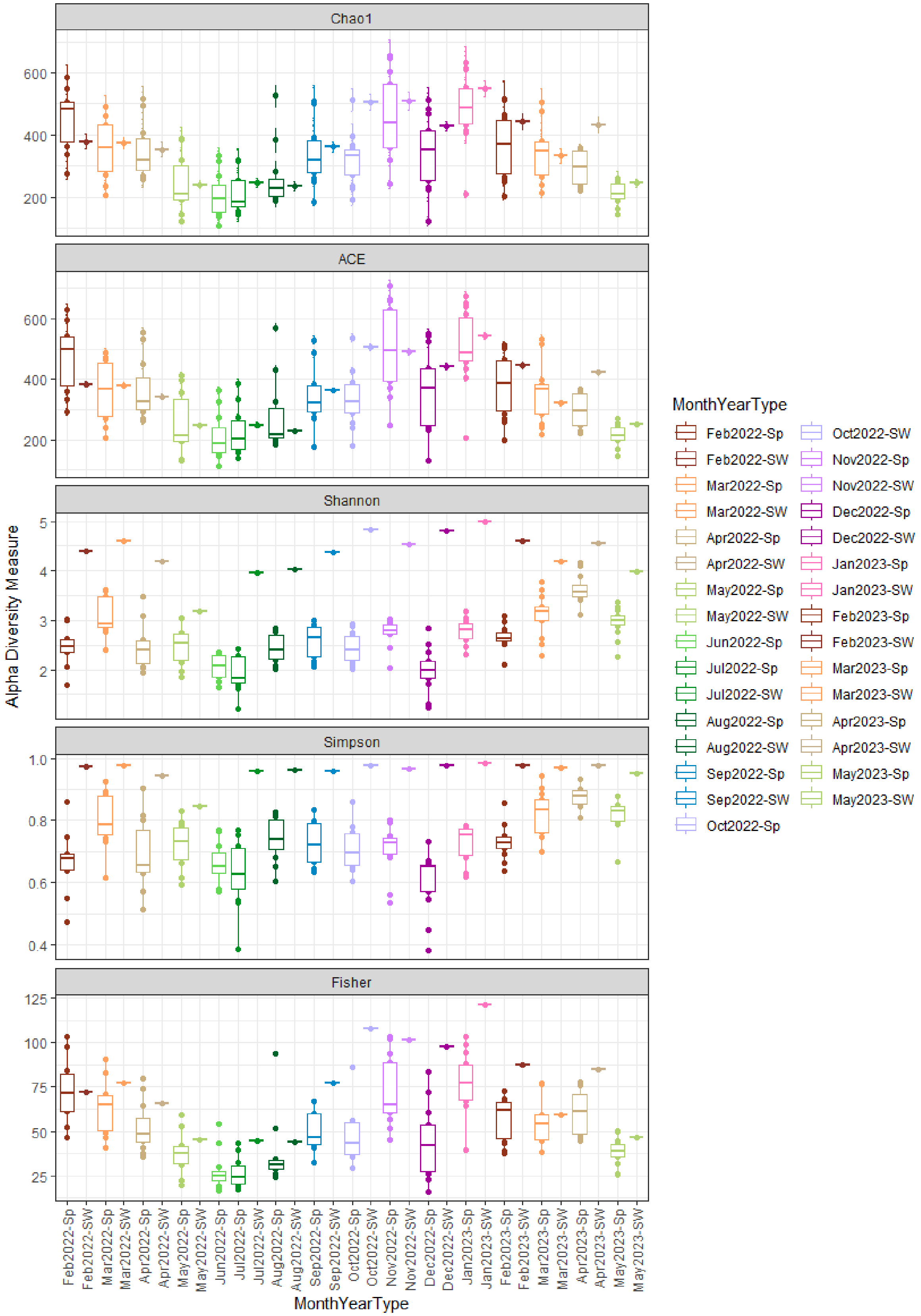


**Figure S6.** Bacterial community diversity and richness in sponge and seawater (SW) over 16 months. Different diversity indices shown as Chao1, ACE, Shannon, Simpson, and Fisher's Alpha.

### Figure S7: NMDS beta diversity Sponge-SW

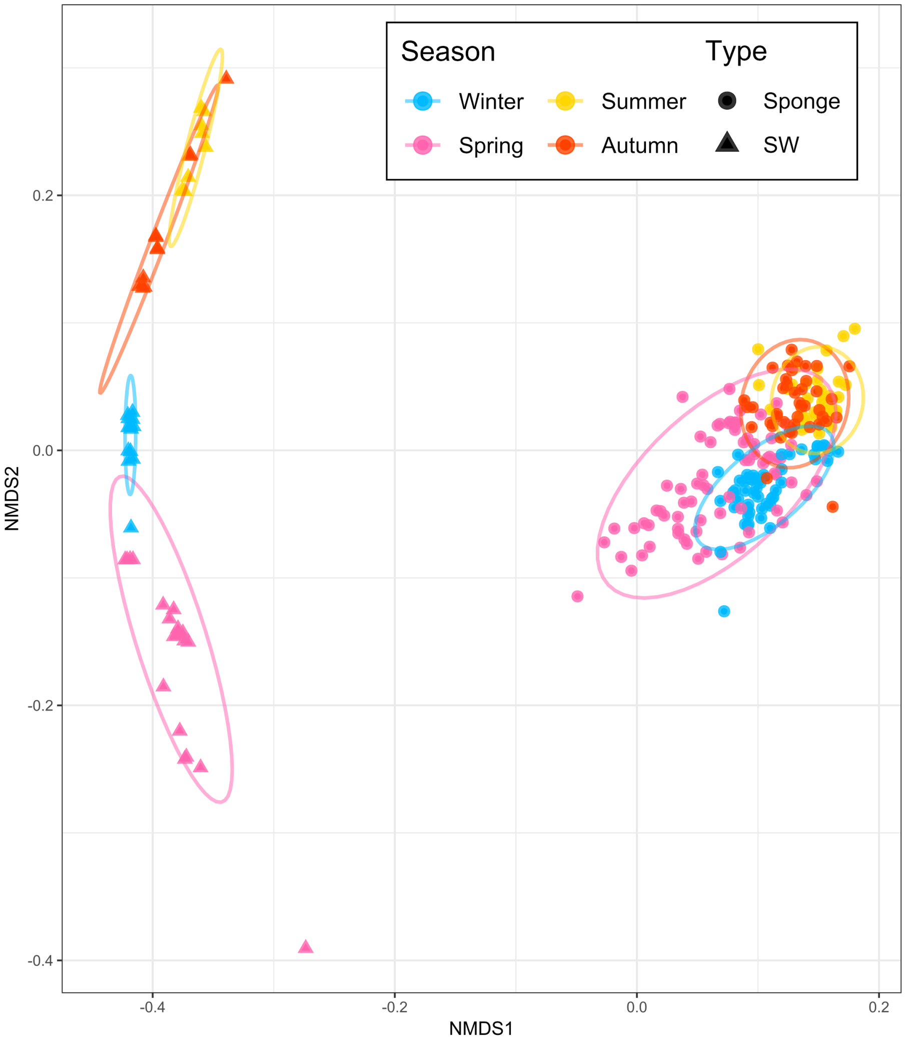


**Figure S7.** NMDS plot showing separation of seawater and sponge samples based on JSD beta diversity, colored according to seasons (stress: 0.093). Ellipses show 95% confidence intervals around group centroids.

### Figure S8: NMDS beta diversity sponge seasons

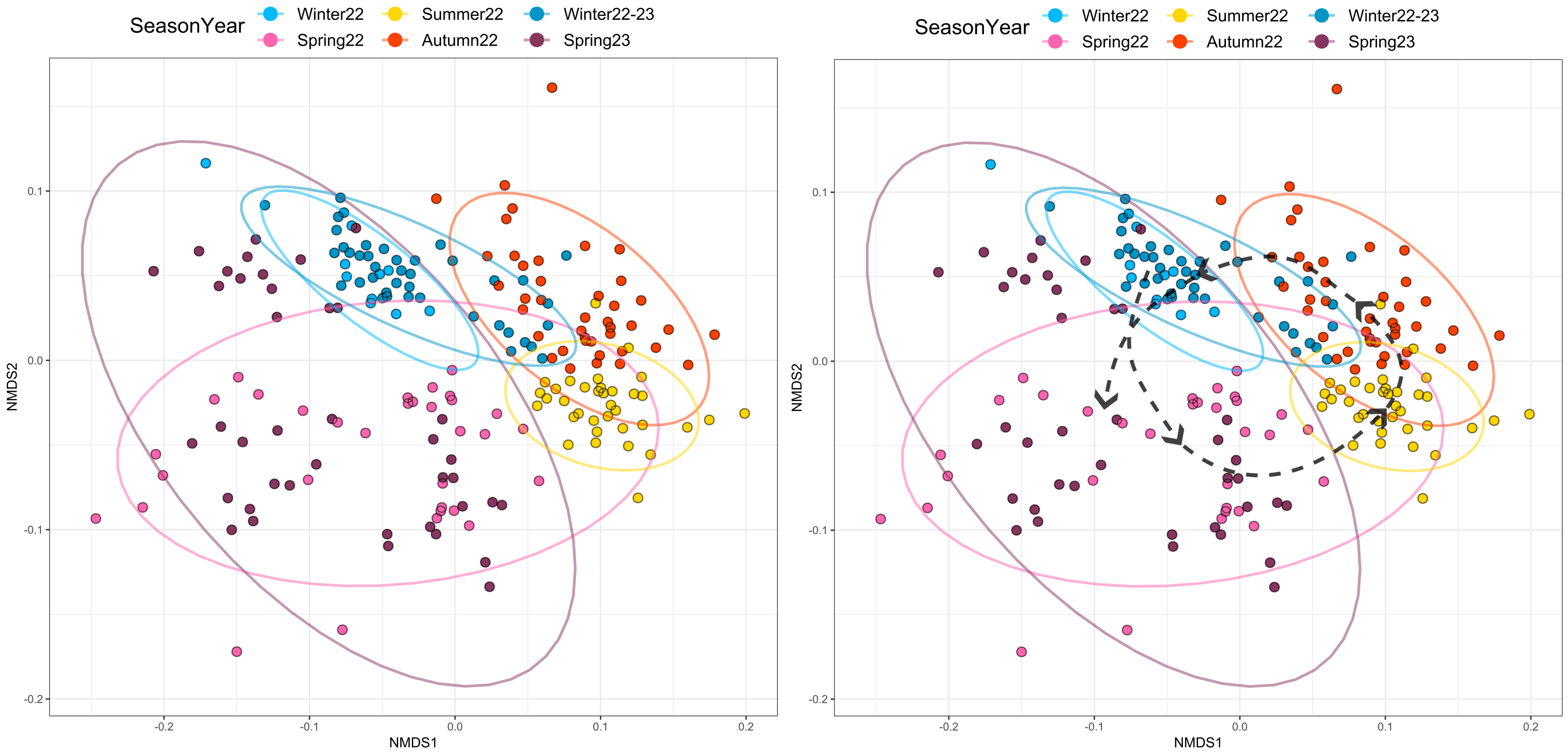


**Figure S8.** NMDS plot showing separation of seasons in sponge samples based on JSD beta diversity, colored by corresponding season and year (NMDS stress: 0.15). Ellipses show 95% confidence intervals around group centroids.

**Figure S8.** NMDS plot showing separation of seasons in sponge samples based on JSD beta diversity, colored by corresponding season and year (NMDS stress: 0.15). Black dashed lines connect centroids of each group. Ellipses show 95% confidence intervals around group centroids.

### Figure S9: Sponge-SW decontamination

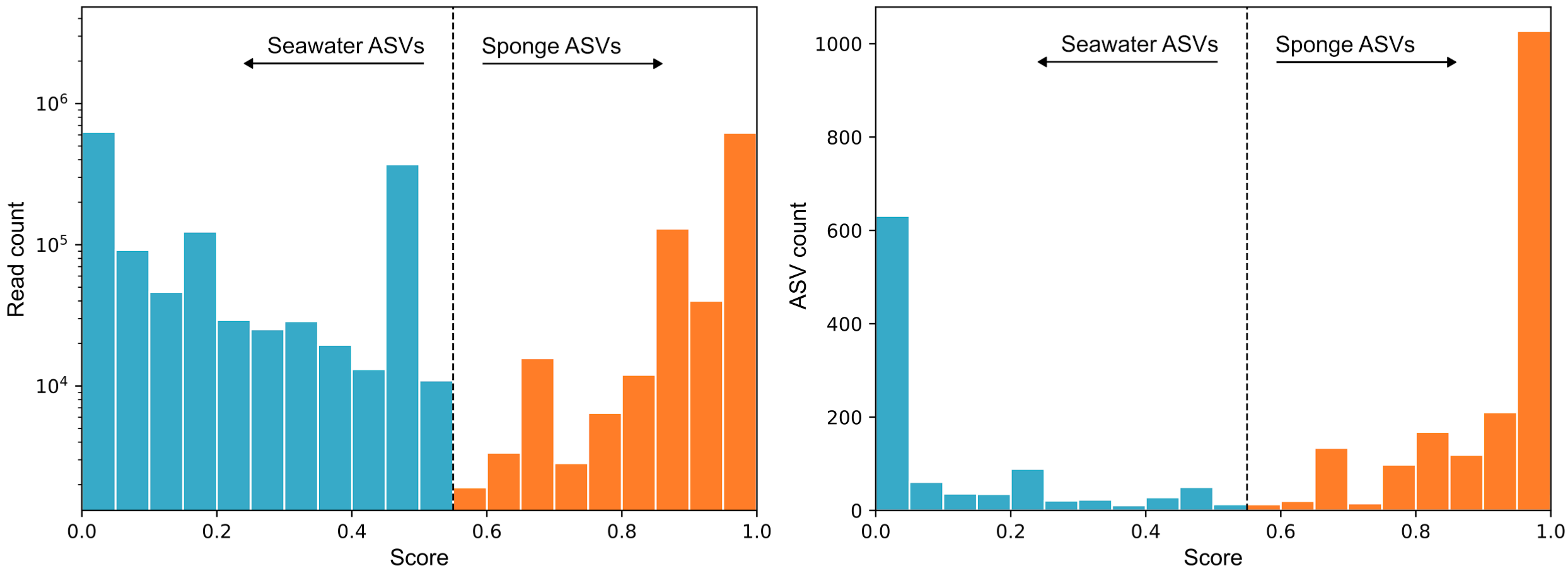


**Figure S9.** Division of ASVs into seawater and sponge groups based on read (left) and ASV (right) frequency distributions. A decontamination score of 0.55 indicated by the dashed line was chosen  as the separation treshold, based on manual inspection of ASV profiles. The prevalence and read abundance of ASVs in seawater samples was screened against all ASVs in sponge samples, and ASVs from the shared pool either remained shared or were relocated to sponge-specific ASVs.

### Figure S10: Core microbiomes features

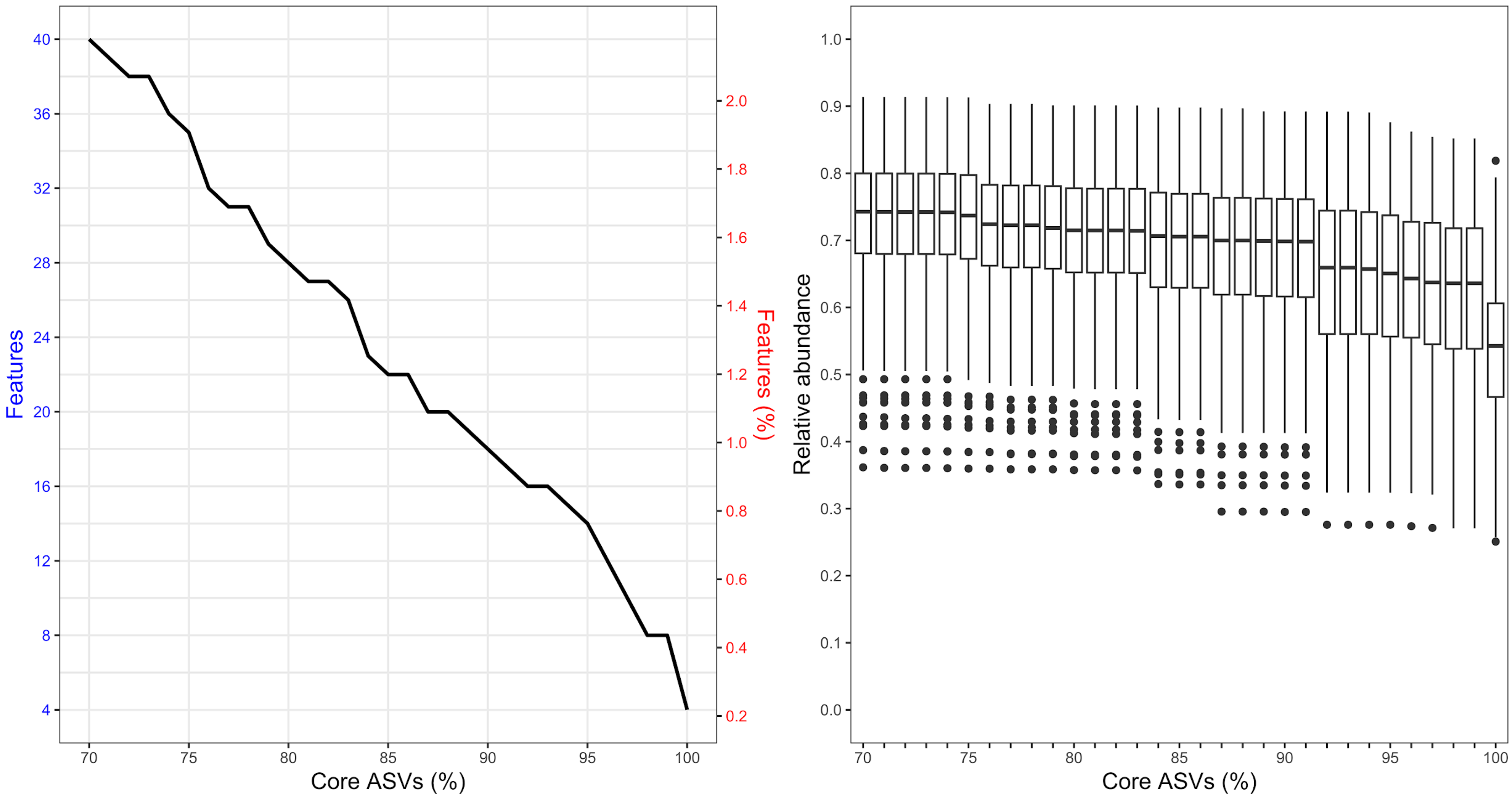


**Figure S10.** Distribution of ASVs (left), and their relative abundance within the microbiome (right), across a range of thresholds (70-100%) for core microbiome detection in sponge samples.
